# Limiting hearing loss in transgenic mouse models

**DOI:** 10.1101/2024.10.03.616327

**Authors:** Travis A. Babola, Naomi Donovan, Sean S. Darcy, Catalina D. Spjut, Patrick O. Kanold

## Abstract

Transgenic mice provide unprecedented access to manipulate and visualize neural circuits, however, those on a C57BL/6 background develop progressive hearing loss, significantly confounding systems-level and behavioral analysis. While outbreeding can limit hearing loss, it introduces strain variability and complicates the generation of complex genotypes. Here, we propose an approach to preserve hearing by crossing transgenic mice with congenic B6.CAST-*Cdh23^Ahl+^*mice, which maintain low-threshold hearing into adulthood. Widefield and two-photon imaging of the auditory cortex revealed that 2.5-month-old C57BL/6 mice exhibit elevated thresholds to high frequency tones and widespread cortical reorganization, with most neurons responding best to lower frequencies. In contrast, *Ahl+* C57BL/6 mice exhibited robust neural responses across a range of frequencies and sound levels (4-64 kHz, 30-90 dB SPL) and retained low thresholds into adulthood. Our approach offers a cost-effective solution for generating complex genotypes and facilitates more interpretable systems neuroscience research by eliminating confounding effects from hearing loss.

## INTRODUCTION

Transgenic animal models have revolutionized our understanding of the brain and provided unprecedented access to cell type-specific monitoring and manipulation. However, many commonly used tools for cell-type specific manipulation, such as conditional knockout alleles (International Mouse Knockout Consortium, 2007), fluorescent reporters (Madisen et al., 2010), and calcium indicators (Dana et al., 2014), are maintained in C57BL/6 mice, which exhibit progressive hearing loss starting at P21 (Keithley et al., 2004; Mikaelian et al., 1974; Zhang et al., 2013). This hearing deficit has been linked to a single nucleotide polymorphism (SNP) in *Cdh23* at the *ahl* (<u>a</u>ge-related <u>h</u>earing loss) locus. This gene encodes Cadherin23, a crucial component of the tip-link structure of the stereocilia in inner hair cells of the cochlea responsible for converting sound waves into neural activity (Noben-Trauth et al., 2003). Therefore, when using adult transgenic mice, researchers must remain aware that these mice progressively lose their ability to hear starting as early as postnatal day P(21) (Zhang et al., 2013), acting as a potential confound in many experimental contexts. Progressive hearing loss is clearly problematic for auditory physiology but also poses challenges for experiments that rely on auditory cues to trigger behavior, such as reward-based learning tasks or maternal responses to ultrasonic pup calls, where auditory perception is essential for normal behavioral outcomes.

Over the past decade, several strategies have been developed to mitigate age-related hearing loss in these mouse models. Commonly, transgenic animals are outbred to strains that carry *Cdh23*^Ahl+^ variants, producing F1 litters that are *Ahl+* (autosomal dominant) and carry the resulting transgene (Frisina et al., 2011; Liu & Kanold, 2021; Romero et al., 2020). One disadvantage of this strategy is that behavior and plasticity can be highly dependent on strain (Kim et al., 2017; Ranson et al., 2013, 2013; Sinclair et al., 2017; Sultana et al., 2019), making comparisons across studies difficult. Moreover, this outbreeding strategy works efficiently only when a single transgene is required. Generating complex transgenic animals, such as using a Cre and Cre-dependent reporter to study a particular cell population, becomes increasingly difficult, as all transgenes must be maintained on a single breeder. Researchers must also avoid Cre lines that result in germline recombination such as *CamKII*α*-Cre*, *Emx1-Cre*, *PV-Cre, GFAP-Cre, Foxg1-Cre,* or *Six3-Cre* (Luo et al., 2020), which are commonly used to study neuronal and glial populations. When conditional knockouts are required, this strategy becomes even more challenging, as knockout alleles (or the Cre) must be outbred and made congenic by multiple rounds of backcrossing to achieve a pure genetic background. Achieving the gold standard of 10 or more generations of backcrossing takes 2-3 years if offspring are randomly selected, with modern speed congenic services taking 5 generations, or 1.5 years (Wong, 2002). More recently, the SNP in C57BL/6 mice was corrected in a single generation using CRISPR/Cas9-mediated genome editing (Mianné et al., 2016), but this process requires considerable resources to design, generate, and screen animals, and carries the additional risk of off-target gene editing. These approaches are impractical for most laboratories and a drain of resources on labs that can afford it.

In this study, we demonstrate that using a commercially available congenic strain, B6.CAST-Cdh23^Ahl+^/Kjn, which has already been outbred and backcrossed to C57BL/6 for over 10 generations, allows for the rapid generation of transgenic animals expressing pan-neuronal GCaMP6s that carry *Ahl+*, therefore limiting the effects of progressive hearing loss. These animals can be genotyped using traditional PCR followed by a restriction enzyme digest, foregoing costly Sanger sequencing to detect *ahl* variants. Using widefield and two-photon imaging, we demonstrate that C57BL/6 mice carrying the *Ahl+* allele exhibit neural responses at thresholds similar to those observed in mice outbred to CBA/CaJ, and retain low thresholds at 6 months of age, within the timeframe of most adult behavior and imaging studies. By identifying the advantages of this approach, we hope to enable better, more interpretable, and cost-effective experiments in neuroscience systems research.

## RESULTS

### Outbreeding transgenic mice limits early hearing loss

A majority of transgenic mice used in modern neuroscience studies are maintained on the C57BL/6 background, a strain known to exhibit signs of progressive hearing loss as early as P21 (Mikaelian et al., 1974; Zhang et al., 2013). One such line is *Thy1-GCaMP6s* (referred to here as *ahl* B6, as it carries two copies of the recessive *ahl* allele), which expresses the genetically encoded calcium indicator GCaMP6s across most excitatory neurons, and is widely used to study circuits *in vivo* (Dana et al., 2014). To determine if these mice exhibit symptoms of early hearing loss, we installed cranial windows and performed widefield imaging of the auditory cortex in awake mice (∼2.5 months old, see STAR Methods for details) while presenting pure tones encompassing their hearing range (4-64 kHz in octave steps, 30-90 dB SPL (decibels, sound pressure level) in 20 dB SPL steps; Figure 1A, B). Pure tones resulted in visible fluorescence increases within discrete spatial domains of the auditory cortex; low frequency tones (4 kHz) presented at moderate sound levels (50-70 dB SPL) typically activated 3 major areas, consistent with A1, A2, and AAF functional designations (Liu et al., 2019; Romero et al., 2020), while higher frequencies activated complex spatial patterns with frequencies advancing from posterior to anterior in A1, dorsal to ventral in A2, and posterior to anterior in AAF (Figure 1C). In general, *ahl* B6 mice exhibited robust, low-threshold responses following presentation of 4, 8, and 16 kHz tones (mean ± SD: 40 ± 20 dB SPL, 30 ± 10 dB SPL, and 40 ± 10 dB SPL respectively; Figure 1D-F), consistent with auditory brainstem responses (ABRs; Ison & Allen, 2004; Johnson et al., 2017). However, responses to higher frequency tones (32 and 64 kHz) at low sound levels were undetectable in most (9 out of 10) mice (Figure 1F). Responses to 32 and 64 kHz tones were elevated (mean ± SD: 80 ± 30 dB SPL and 80 ± 20 dB SPL respectively), indicating a loss of hearing sensitivity at these frequencies (Figure 1F). These data indicate that even at a young age, high frequency hearing is significantly impaired.

**Figure 1.**
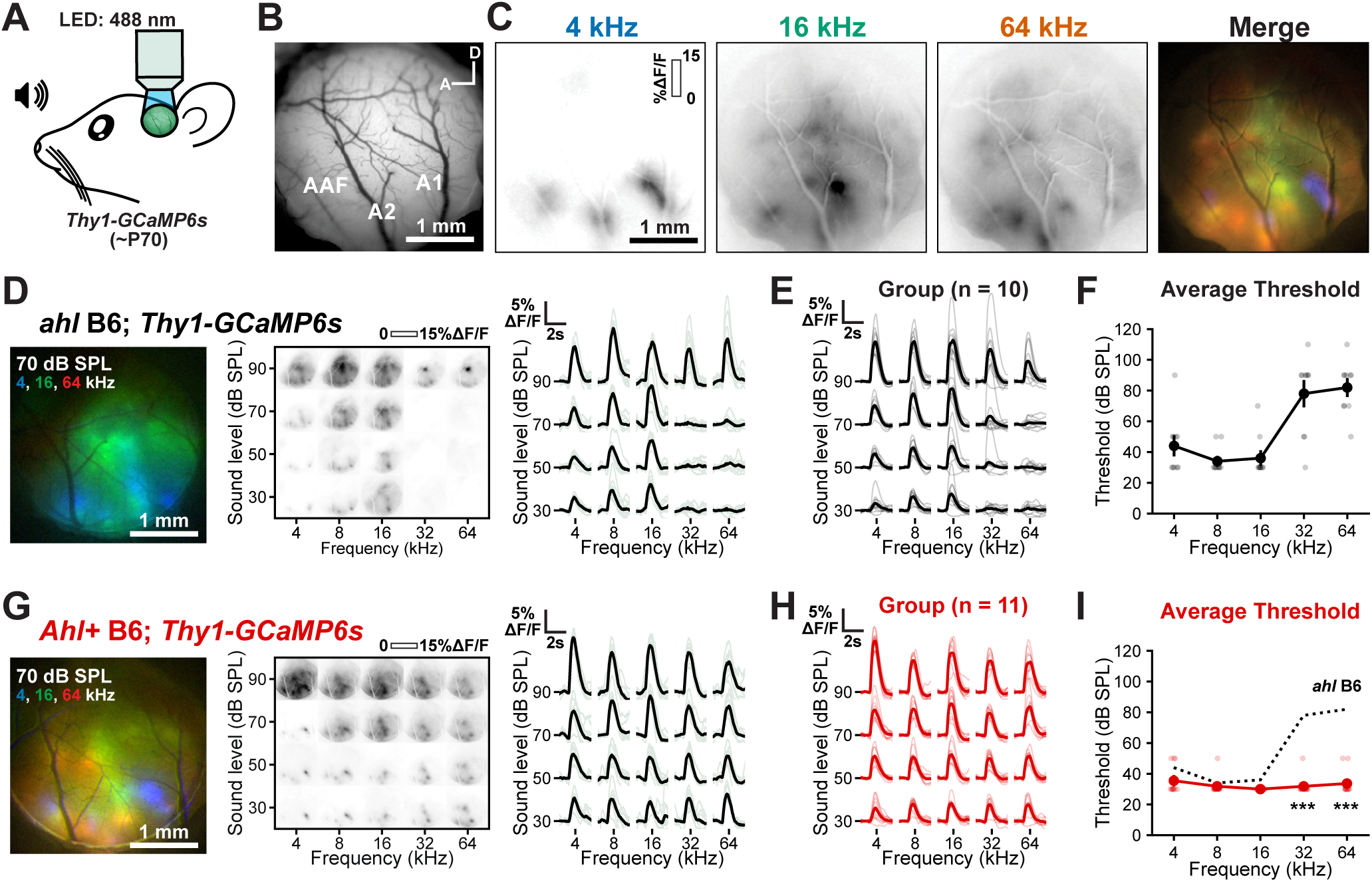
Widefield imaging of sound-evoked responses in the auditory cortex. **A,** Schematic of widefield imaging in auditory cortex of adult (∼P70) *Thy1-GCaMP6s* mice. **B,** Representative image of the cranial window and the field of view captured for analysis. **C,** Fluorescence responses to pure tones indicated at 70 dB SPL. (right) Merged fluorescence responses to pure tones highlight cortical topography. Blue, green, and red channels indicate responses to 4, 16, and 64kHz, respectively. **D,** (left) Merged fluorescence responses to 4 (blue), 16 (green), and 64kHz (red) at 70 dB SPL in *ahl* B6; *Thy1-GCaMP6s* mice. (middle) Plot of fluorescence changes over the imaging field to varying frequency (x-axis) and sound levels (y-axis). (right) Plot of fluorescence during sound presentation from an individual mouse; grey lines are individual trials, black line is the average. **E,** Plot of fluorescence during sound presentation from all *ahl* B6; *Thy1-GCaMP6s* mice; grey are individual mice, black is the group average, n = 10 mice. **F,** Plot of average fluorescence detection threshold for each frequency presented. Light individual markers represent individual mice. If there was no observable response at 90 dB SPL, the threshold was defined as 110 dB SPL. **G-I**, Similar to D-F, but for *Ahl+* B6*; Thy1-GCaMP6s* mice. Three-way ANOVA (threshold, sex, and genotype reported in text), followed by post-hoc t-tests with Benjamini-Hochberg FDR correction. ***p<0.001.

To limit hearing loss in transgenic mice, a common strategy is to outbreed them to a strain that carries the *Ahl+* allele, such as CBA/CaJ (Bowen et al., 2020; Liu et al., 2019). Indeed, we observed robust widefield calcium responses to all frequencies and attenuations tested in F1 offspring using this breeding strategy ((CBA/Ca x C57BL/6J)F1, referred to here as *Ahl+* B6.CBA; Supp Figure 1). While this approach is tractable in animals carrying a single allele, generating complex transgenic animals, such as conditional knockouts, is impossible while maintaining a consistent genetic background without generating congenic animals.

To bypass these difficulties, we used a commercially available congenic strain, *B6.CAST-Cdh23^Ahl+^/Kjn*, which has already been outbred and backcrossed to C57BL/6J for over 10 generations while selecting for the *Ahl+* allele. Similar to *Ahl+* B6.CBA mice, breeding *Thy1-GCaMP6s* mice to *B6.CAST-Cdh23^Ahl+^/Kjn* mice produced offspring (*Ahl+* B6) that exhibited neural responses to tones across all frequencies and attenuations tested (Figure 1G, H). To compare auditory thresholds across all frequencies, genotypes, and sexes, we performed a three-way ANOVA, which revealed significant effects for all main factors: frequency: (F(4,110) = 16.2, *p* < 0.001), sex (F(1,110)=4.75, *p* = 0.031), and genotype (F(2,110) = 61.7, *p* < 0.001), as well as their interactions (F(22,110)=5.1, *p* < 0.001). Although sex was a weakly significant factor, post-hoc testing revealed no statistically significant differences between sexes within each genotype (t-tests with corrected p-value, *Ahl+* B6, *p* = 0.097; *Ahl+* B6.CBA, *p* = 0.462; *ahl* B6*, p* = 0.53). In contrast, frequency and genotype were very strong predictors. The *ahl* B6 mice exhibited higher thresholds at 32 and 64 kHz compared to both *Ahl+* B6 (mean ± SD: 32 ± 5 and 34 ± 8 dB SPL for 32 and 64 kHz, t-test with corrected *p* < 0.001 and *p* < 0.001, respectively) or *Ahl+* B6.CBA mice (30 ± 0 and 33 ± 8 dB SPL for 32 and 64 kHz, t-tests with corrected *p* = 0.001 and *p* < 0.001, respectively) (Figure 1I and Supp Figure 1C). These data indicate that the *Ahl+* locus is critical for preserving high frequency hearing, consistent with previous findings (Johnson et al., 2017). By implementing a simple cross to B6.CAST*-Cdh23^Ahl+^*/Kjn, our strategy enables the rapid generation of complex transgenic mice with limited hearing loss.

### c.573A variant introduces a restriction enzyme site that allows for fast genotyping without sequencing

To maintain and generate new breeders and offspring carrying the *Ahl+* allele, we developed a rapid genotyping method. Typically, detecting SNPs like the c.573A variant (*ahl*) in C57BL/6 mice involves amplifying the region with PCR and subsequently submitting the sample for sequencing (Figure 2A). However, the B6 *ahl* variant introduces a BsrI restriction enzyme site (Figure 2B). By offsetting the PCR primers (Figure 2A), we found that amplifying an ∼950 base pair region of *Cdh23* (exon 9) and then digesting with BsrI can differentiate between mice with zero, one, or two copies of the recessive *ahl* allele. In *Ahl+* homozygous mice (*Cdh23*^Ahl/Ahl^), this method produces a single, uncut band of 948 base pairs (Figure 2C). In *ahl* homozygous mice (*Cdh23*^ahl/ahl^), the DNA is cut by BsrI, resulting in bands at 284 and 664 base pairs. Heterozygous mice exhibit all three bands (Figure 2C). This procedure takes approximately two hours, does not require DNA purification, and can be performed with a standard thermocycler (see Materials and Methods), offering a quick, efficient, and accessible alternative to sequencing or outsourcing to distinguish *Cdh23* variants.

**Figure 2.**
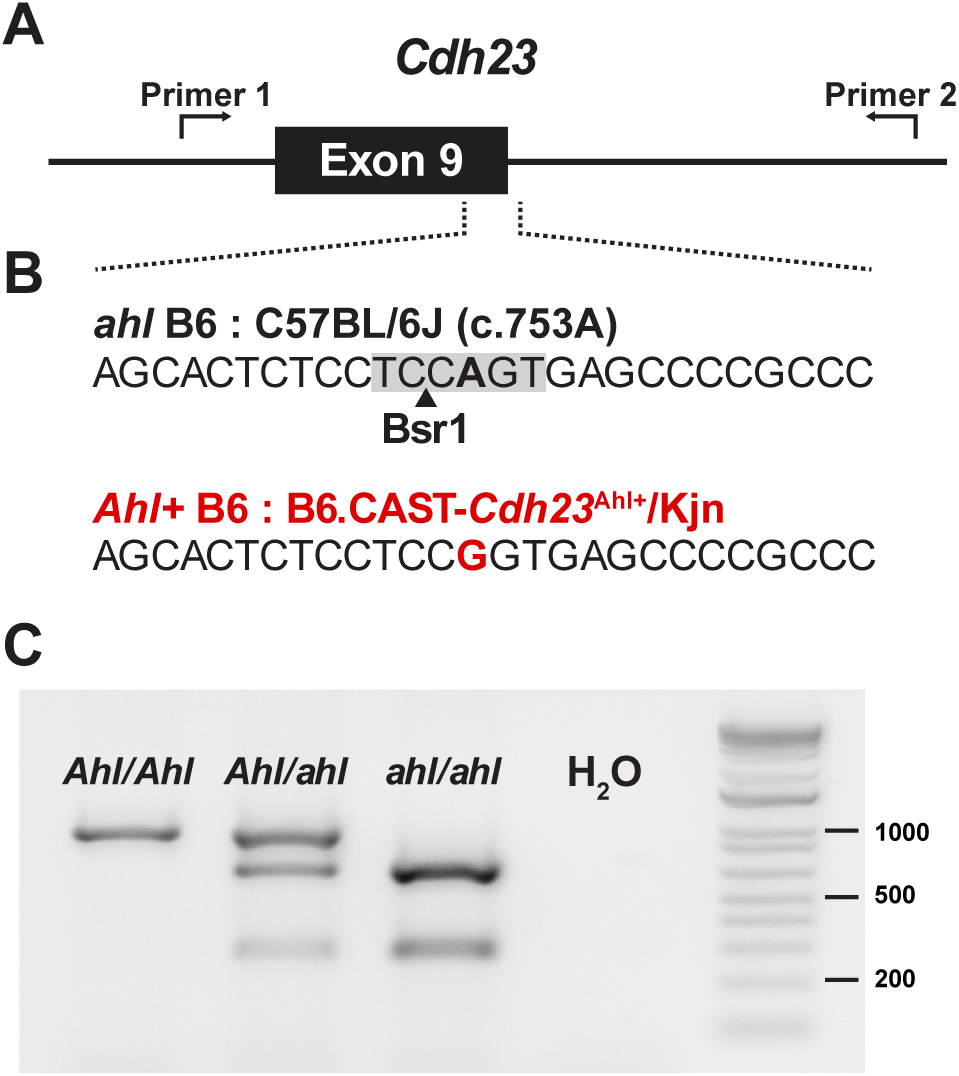
Genotyping *Cdh23* SNP responsible for hearing loss in C57BL/6 mice. **A,** Schematic of *Cdh23* locus (exon 9), where C57BL/6 mice exhibit a SNP. Primers are designed such that the SNP is offset from the center of the amplified region. **B,** Sequence of the region denoted in A from C57BL/6 (top) and B6.CAST-*Cdh23*^Ahl+/^Kjn (bottom) mice. A BsrI restriction enzyme site is present in C57BL/6. **C,** Image of gel following PCR amplification and BsrI restriction enzyme digest of DNA region shown in **A** in mice carrying zero, one, or two *ahl* alleles.

### Two-photon imaging reveals diminished responses to high frequency tones in *ahl* B6 mice

To define the response properties of individual neurons, we performed two-photon imaging within the auditory cortex in awake mice. Using the tonotopic maps identified with widefield imaging as a reference (Figure 3A), the imaging field of view encompassed the primary auditory cortex (A1; ∼1.1mm^2^). We imaged 250 µm below the pial surface, anatomically at the bottom of layer III and the top of layer IV (de Vries et al., 2020). Neurons within A1 exhibited spatially organized responses to tones (Figure 3A), with many that were broadly responsive to louder tones and selective to quieter tones (Figure 3B), characteristic of “V-shaped” tuning. Overall, there were no significant differences in the number of sound-responsive neurons in *Ahl+* B6 or B6.CBA mice compared to *ahl* B6 mice (Figure 3C and Supp Figure 2A). However, the proportions of neurons responding to sound offset were significantly higher in *Ahl+* B6 and B6.CBA mice compared to *ahl* B6 mice (Supp Figure 3A), similar to *Ahl+* B6.CBA mice in previous studies (Bowen et al., 2020). When examining the average tuning curve across all sound-responsive neurons, neuronal response properties were remarkably consistent with widefield imaging. The *ahl* B6 mice lacked responses to high frequency tones (32 and 64 kHz) at lower sound levels, while *Ahl+* B6 mice exhibited responsiveness across all frequencies and sound levels (Figure 3D). These data indicate that *ahl* B6 mice lack neurons responding to high frequencies across A1 and that widefield imaging serves as a reasonable proxy for assessing general cortical responsiveness to sounds.

**Figure 3.**
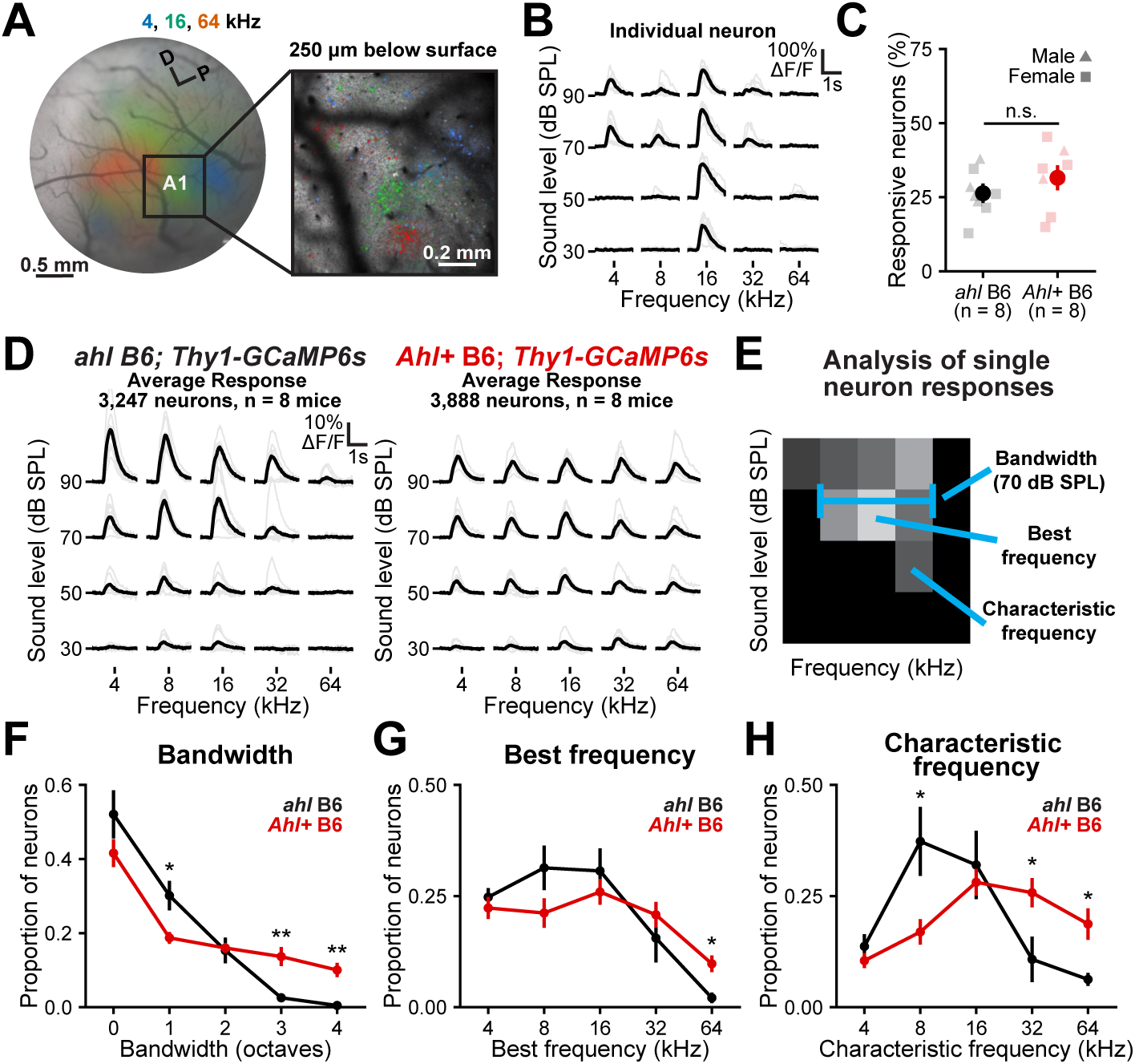
Two-photon imaging of sound-evoked responses in auditory cortex. **A,** (left) Schematic of widefield imaging in auditory cortex of adult (P60) *Ahl+* B6; *Thy1-GCaMP6s* mice with overlaid responses to 70 dB SPL tones. (right) Two-photon imaging of primary auditory cortex (A1, 250 µm from pial surface) with overlaid responses to 70 dB SPL tones. **B,** Fluorescence traces from an individual neuron within the field of view depicted in **A** as a function of frequency (x-axis) and sound level (y-axis). Grey traces are individual trials, black traces are the trial averages. **C,** Plot of the proportion of sound-responsive neurons within each genotype. Light markers indicate individuals and dark markers are mean ± SEM, n = 8 mice for each genotype. Two-way ANOVA, genotype: *F(2,15) =* 2.42, *p* = 0.12; sex: *F(1,15)* = 1.95*, p* = 0.18, interaction: *F(2,15)*=0.00*, p* = 0.99. n.s.: not significant. **D,** Average response across all sound-responsive neurons. Grey traces are individual mice, black traces are the average. n = 8 mice for each genotype, 3,247 total neurons (mean ± SD: 406 ± 190) in *ahl* B6 mice, 3,888 total neurons (mean ± SD: 486 ± 258) in *Ahl+* B6 mice **E,** Schematic indicating measurements reported in **F-H.** Color block indicates the response of an individual neuron to different frequencies and sound levels, with lighter colors indicating larger responses. Bandwidth is defined as the width in octaves separating the highest and lowest responding frequencies; a bandwidth of 0 indicates the neuron was responsive to a single tone. Best frequency is defined as the frequency with the highest amplitude response, regardless of sound level. Characteristic frequency is defined as the frequency eliciting the highest amplitude response at the lowest sound level. **F-H,** Plots of the proportion of neurons as a function of bandwidth at 70dB, best frequency, and characteristic frequencies across genotypes. Markers are mean ± SEM, n = 8 mice for each genotype. Two-way ANOVAs with genotype and measurement (see Supp Statistics for values), followed by post-hoc t-tests with Benjamini-Hochberg FDR correction. *: *p* < 0.05, **: *p* < 0.01, ***: *p* < 0.001.

To assess how neurons in a network connect and organize, we calculated signal correlations, which measure the similarity in sound-evoked responses between pairs of sound-responsive neurons, and noise correlations, which reflect the similarity in activity that is independent of the stimuli. When averaging across all frequency and attenuation levels, we observed no significant differences between *Ahl+* B6 and *ahl* B6 mice in either signal or noise correlations (Supp Figure 3B-C). However, *Ahl+* B6.CBA mice exhibited lower signal correlations than *Ahl+* or *ahl* B6 mice (Supp Figure 3B), suggesting a sparser representation of tones in these mice (Bowen et al., 2020).

To understand how the response properties of individual neurons differ between genotypes, we calculated standard tuning properties for each neuron. The proportion of neurons with wide bandwidths, defined as the range in octaves between the minimum and maximum responsive frequency at 70 dB SPL (Figure 3E), was higher in *Ahl+* B6 mice than in *ahl* B6 (Figure 3F), consistent with the lack of response to 32 and 64 kHz tones in *ahl* B6 mice at this sound level (Figure 3D). Similarly, *Ahl+* B6 mice exhibited higher proportions of neurons that responded with the highest amplitude to 64 kHz, regardless of sound level (best frequency, Figure 3G). Remarkably, when examining the characteristic frequency, defined as the frequency that elicited the highest response at the lowest sound level, neurons in *ahl* B6 mice exhibited a dramatic shift toward lower frequencies, with a significantly higher proportion of neurons with characteristic frequencies of 8 kHz and significantly lower proportions of neurons with characteristic frequencies of 32 and 64 kHz (Figure 3H). Given similar numbers of sound-responsive neurons across genotypes (Figure 3C), these data suggest that connections among neurons within *ahl* B6 mice had reorganized, allocating more cortical area to lower frequencies and reflecting the concurrent loss of high frequency sensitivity in the inner ear.

### Reorganization of cortical responses towards lower frequencies in *ahl* B6 mice

To directly assess if reorganization of response properties occurred across the spatial extent of A1, we generated spatial characteristic frequency maps that mark the location and characteristic frequency of each neuron (Figure 4A). Using a winner-takes-all approach, we summed the responses of local neurons exhibiting the same characteristic frequency using a Gaussian filter (σ = 60 µm) and assigned the frequency with the highest value to that unit of area (Figure 4B). Compared to *Ahl+* B6 and *Ahl+* B6.CBA mice, which exhibited frequencies distributed across the field of view (Figure 4C, Supp Figure 2C, D), *ahl* B6 mice displayed a remarkable shift towards processing lower frequencies (≤16 kHz), with a majority (86 ± 14%) of the cortical area responding best to that range. We then reexamined signal correlation measurements by conditioning the correlations on lower frequencies and higher sound levels, where *ahl* B6 mice are most likely to respond, and found that signal correlations were much higher (r mean ± STD: 0.16 ± 0.04) in *ahl* B6 mice compared to *Ahl+* B6 (0.10 ± 0.02; t-test with corrected *p* = 0.004) and B6.CBA (0.03 ± 0.01; t-test with corrected *p* < 0.001) mice (Supp Figure 3D), consistent with a reorganization towards lower frequencies. This increase in signal correlations likely reflects increasing amounts of shared input from thalamocortical pathways or strengthened synaptic strength of spared inputs. We did not observe cortical areas devoid of auditory responses, which might be predicted after a sudden loss of high frequency hearing (Noreña et al., 2010), suggesting that auditory circuits had already undergone plasticity and/or increased cortical gain in response to degraded high-frequency input (Chambers et al., 2016; McGill et al., 2022; Willott et al., 1993).

**Figure 4.**
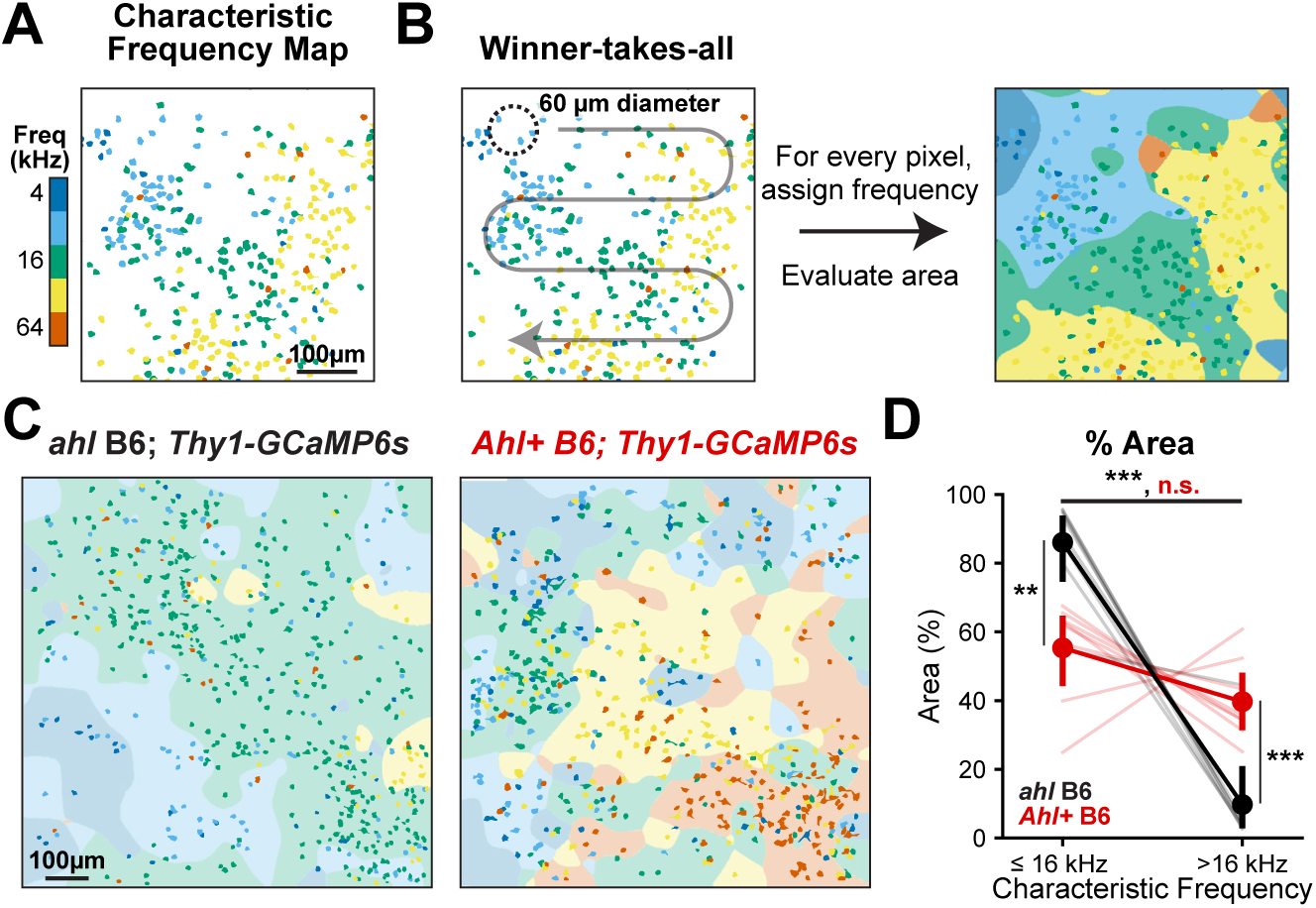
Reorganization of auditory cortex towards lower frequencies in *ahl* B6 mice. **A,** Plot of neuron location with characteristic frequencies indicated with color. **B,** Schematic of the analytic method to quantify cortical area devoted to processing a given characteristic frequency. A Gaussian filter (σ = 60 µm) was used to create a weighted sum of each neuron’s characteristic frequency response for each unit of area. The frequency with the highest value was assigned to that unit of area and proportion of area examined. **C,** Exemplar characteristic frequency maps and assigned areas for the indicated genotypes. **D,** Plot of the fractional area as a function of characteristic frequency range and genotype. Light lines are individual animals, dark lines are mean ± SEM, n = 8 mice for each genotype. Two-way ANOVA (characteristic frequency: *F(1,36)* = 72.98*, p* < 0.001; genotype: *F(2,36) = 0.08, p* = 0.92; interaction: *F(2,36)* = 34.6, *p* < 0.001), followed by post-hoc t-tests with Benjamini-Hochberg FDR correction. n.s: not significant, **: *p* < 0.01, ***: *p* < 0.001.

### Linear predictive model of population responses performs worse at high frequencies and low sound levels in *ahl* B6 mice

C57BL/6 mice are commonly used in behavioral experiments involving acoustic stimuli (Breton-Provencher et al., 2022; Guo et al., 2019; Inagaki et al., 2018), but our results suggest that these animals process sounds differently. While responses to high frequencies are much lower in *ahl* B6 mice, there remains a small population of neurons that do respond (Figure 3G, H), suggesting that the cortex could decode those tones, such as in the case of mothers responding to pup calls (Marlin et al., 2015). To understand which frequencies and sound levels contain the most information, we designed a linear classifier to predict both the frequency and sound level being presented given linear features extracted from the population of neuronal responses (Figure 5A). The population responses were first transformed into a lower dimensional space using principal component analysis, before training a linear discriminate analysis model with 10-fold cross-validation. In *ahl* B6 mice, the population response contained less information relative to *Ahl+* B6 mice, as indicated by overlapping trial responses in lower dimensional space and more incorrect predictions with 32 and 64kHz tone classification (Figure 5B). In contrast, *Ahl+* B6 and B6.CBA mice had well-separated clusters and a high percentage of correct predictions across frequency and sound levels (Figure 5C, Supp Figure 2E). Overall, the linear classifier performed significantly better at 4, 16, 32, and 64 kHz in *Ahl+* B6 mice when combining all attenuation levels (Figure 5D). Examining accuracy as a function of sound level, both models performed similarly at lower frequencies and above chance (5%) at higher frequencies at 90 dB SPL (Figure 5E). At 50 and 70 dB SPL, accuracy at 32 and 64 kHz was much lower in *ahl* B6, but slightly above chance (Figure 5E), indicating preservation of information at these sound levels. Information at 8 and 16 kHz was largely preserved across all sound levels across mice, with no statistical significance between the groups between 30-90 dB SPL (Figure 5E). Similarly, *ahl* B6 mice exhibit highly predictive neural responses to 4 kHz at 70 and 90 dB SPL, with less predictive power at lower sound levels compared to *Ahl+* B6 mice, consistent with weak responses observed with widefield and two-photon imaging (Figure 1E and 3D). Taken together, these data suggest that loud sounds can evoke discriminative properties in population neuronal responses in *ahl* B6 mice, while softer sounds cannot. Therefore, behavioral experiments using *ahl* B6 mice could be confounded by diminished peripheral and cortical responses to low sound levels.

**Figure 5.**
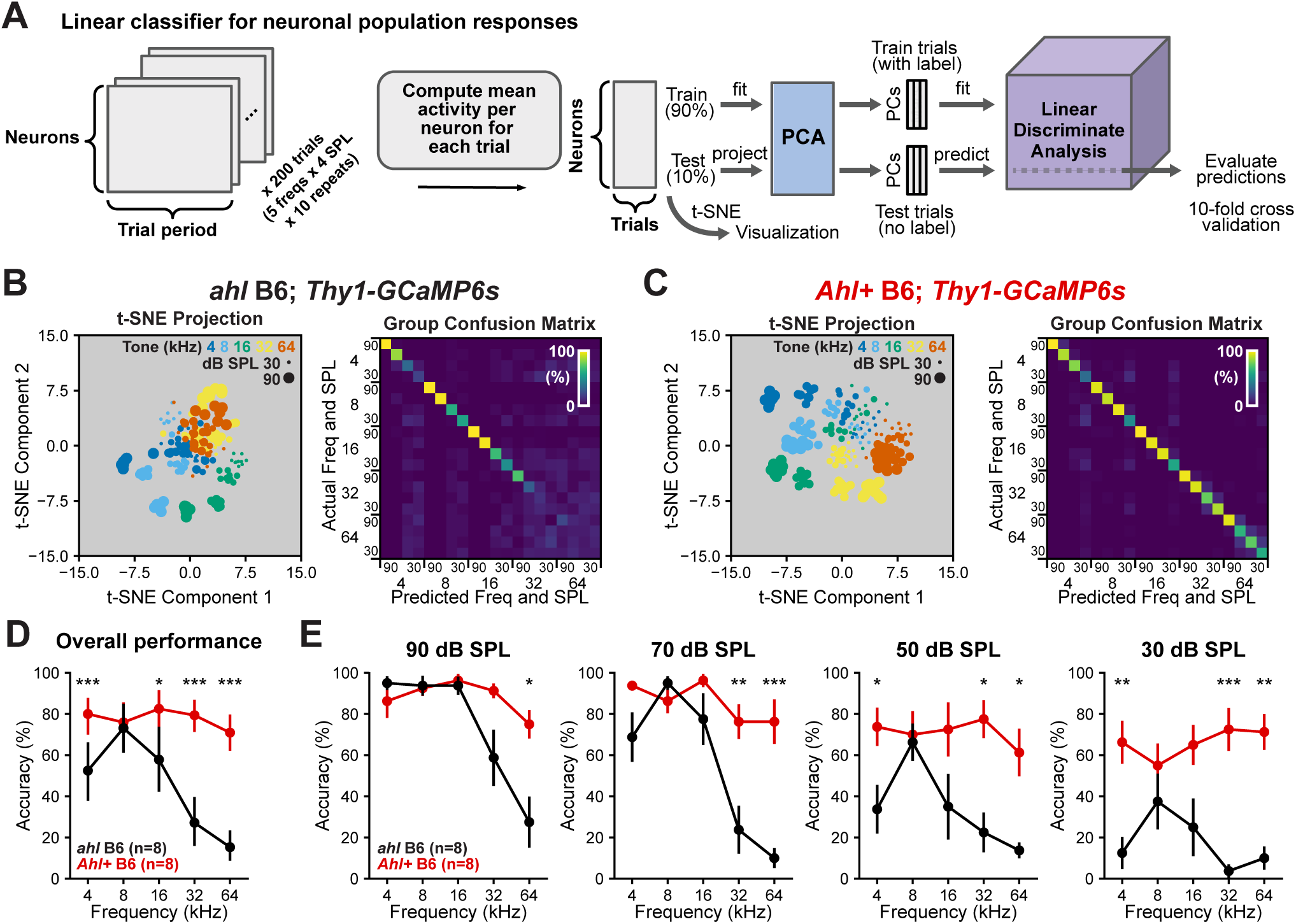
Linear classifier of network response performs worse in *ahl* B6 mice. **A,** Schematic of a pipeline for prediction of frequency and sound level based on population-level response using linear discriminant analysis. **B-C,** (left) Low-dimensional representation (t-SNE) of neuronal population response from an individual animal with each marker representing a single trial. (right) Confusion matrix of classifier performance from a single animal. **D,** Plot of overall classifier performance as a function of frequency. Markers are mean ± SEM, n = 8 mice for each genotype. Three-way ANOVA (genotype: *F(2,360)* = 76.7*, p* < 0.001; frequency: *F(4,360)*=15.2*, p* < 0.001; sound level: *F(3,360)* = 47.8*, p* < 0.001; interaction: *F(50,360)*=2.89*, p* < 0.001), all attenuation levels included for each frequency in post-hoc t-tests with Benjamini-Hochberg FDR correction. *: *p* < 0.01, **: *p* < 0.01, ***: *p* < 0.001. **E,** Plot of classifier performance as a function of frequency (x-axis) and sound level (plots arranged from highest sound level to lowest sound level). Markers are mean ± SEM, n = 8 mice for each genotype. The same three-way ANOVA as reported in **D,** post-hoc t-tests with Benjamini-Hochberg FDR correction. *: *p* < 0.01, **: *p* < 0.01, ***: *p* < 0.001.

### *Ahl+* B6 mice retain hearing sensitivity at 6 months of age, while *ahl* B6 mice lose it

Researchers must invest significant time to train mice on behavioral tasks. Given the progressive nature of hearing loss in C57BL/6 mice, mice could experience varying hearing sensitivity over the training period. To quantify the degree of this hearing loss, we performed widefield imaging on the same cohort of mice at 6 months of age (Figure 6A, B). In *ahl* B6 mice, there was a near-total loss of sensitivity to tones presented at both 30 and 50 dB SPL (Figure 6B). The thresholds for detecting a sound-evoked response significantly increased across all frequencies tested (ANOVA followed by t-tests with corrected p-values: *p* < 0.001, *p* < 0.001, *p* = 0.002, *p* = 0.007, *p* = 0.012 for 4, 8, 16, 32, and 64 kHz respectively), with most mice exhibiting no sound-evoked response for 32 or 64 kHz tones at the highest sound level presented (90 dB, Figure 6B, C). *Ahl+* B6 and B6.CBA mice retained sound-evoked responses across the entire frequency and attenuation range tested, with no significant differences observed in sound-evoked response thresholds (Figure 6D-F and Supp Figure 1D-F). Moreover, 6-month-old *Ahl+* B6 and B6.CBA thresholds were no different or lower than 2.5-month-old *ahl* B6 mice (Figure 6F, Supp Figure 1F). While *Ahl+* B6 mice exhibited 6-month thresholds that were not statistically significant from 2.5-month thresholds, there was a trend towards less sensitivity at the lowest and highest frequencies tested (Figure 6F). *Ahl+* B6.CBA mice do not show this same loss of sensitivity (Supp Figure 1F), consistent with ABR studies that indicate that *Ahl+* B6 mice have slightly elevated hearing thresholds compared to outbred mice (Kane et al., 2012). Together, these data indicate that *Ahl+* B6 mice retain and *ahl* B6 lose a majority of their hearing sensitivity in the first 6 months of life.

**Figure 6.**
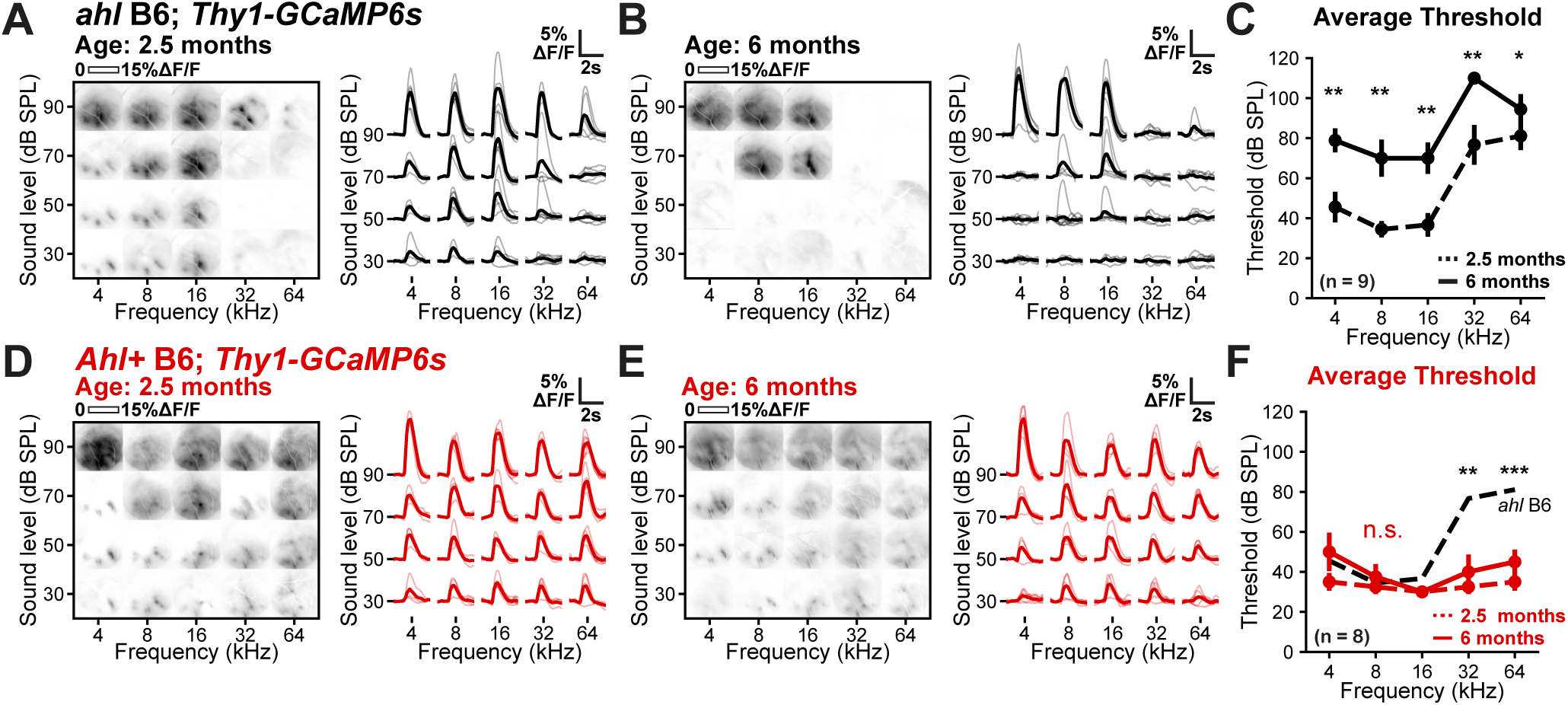
Low threshold responses are intact in 6-month-old *Ahl+* B6 mice. **A,** (left) Plot of fluorescence changes in 2.5-month-old *ahl* B6 mice over the imaging field to varying frequency (x-axis) and sound levels (y-axis). (right) Plot of fluorescence during sound presentation across animals; grey traces are individual mice, black traces are the average, n = 9 mice. **B,** Plot of fluorescence changes in 6-month-old *ahl* B6 mice over the imaging field to varying frequency (x-axis) and sound levels (y-axis). (right) Plot of fluorescence during sound presentation across animals; grey traces are individual mice, black is the average, n = 9 mice. Note the complete disappearance of responses to 32kHz and to 4-16kHz at low sound levels. **C,** Plot of average threshold as a function of frequency and time point. Dashed lines indicate measurements at 2.5 months, solid lines indicate measurements at 6 months, n = 9 mice. Markers at 110 dB SPL indicate no response was observed at 90 dB SPL. Three-way ANOVA (frequency: *F(4,70)* = 10.6*, p* < 0.001; timepoint: *F(1,70)* = 97.8*, p* < 0.001; sex: *F(1,70)* = 0.29*, p* = 0.59, interaction: *F(13,70)* = 1.51*, p* = 0.14) with post-hoc paired t-tests with Benjamini-Hochberg FDR correction. *: *p* < 0.01, **: *p* < 0.01, ***: *p* < 0.001. **D,** Similar to A, but for *Ahl+* B6 mice. n = 8 mice. **E,** Similar to B, but for *Ahl+* B6 mice. n = 8 mice. **F,** Similar to C, but for *Ahl+* B6 mice. n = 8 mice. Three-way ANOVA (frequency: *F(4,60)* = 2.26*, p* = 0.07; timepoint: *F(1,60)* =7.83*, p* = 0.007; sex: *F(1,60)*=0.29*, p* = 0.009, interaction: *F(13,60)* = 0.71*, p* = 0.74) with post-hoc paired t-tests with Benjamini-Hochberg FDR correction. n.s.: not significant.

## DISCUSSION

Transgenic mice are transformative tools that allow precise manipulation and visualization of neural circuits. Because many commonly used transgenic mice are maintained on a C57BL/6 genetic background (Dana et al., 2014; de Vries et al., 2020), they exhibit progressive hearing loss (Mikaelian et al., 1974), acting as a potential confound in studies examining many aspects of brain function. Common strategies used to circumvent this deficit, such as generating CRISPR/Cas9 single nucleotide variants or congenic mice, are time- and resource-intensive and simply not feasible for most laboratories.

Here, we highlight a simple and cost-effective strategy to generate transgenic C57BL/6 mice with limited hearing loss. By crossing commercially available congenic B6.CAST-*Cdh23*^Ahl+^/Kjn mice to pan-neuronal *Thy1-GCaMP6s* mice, we generated offspring with the *Ahl+* allelic variant, known to limit progressive hearing loss (Keithley et al., 2004). These mice exhibited low thresholds to high frequency tones and retained these thresholds at 6 months of age, similar to mice outbred to the CBA/CaJ strain (Figure 6, Supp Figure 1). In contrast, mice without this variant (C57BL/6) exhibited elevated thresholds to high frequencies in early adulthood (Figure 1, 3), reorganization of auditory cortex to respond best to low frequency tones (Figure 4), and progressive loss of sensitivity in the first half-year of life (Figure 6). We show that this variant is easy to genotype with traditional PCR and restriction enzyme digest, preventing the need for DNA purification and sequencing (Figure 2). This strategy is scalable to more complex genetic strategies, such as conditional knockouts or other models requiring multiple transgenes (Figure 7), enabling more interpretable studies in neuroscience.

**Figure 7.**
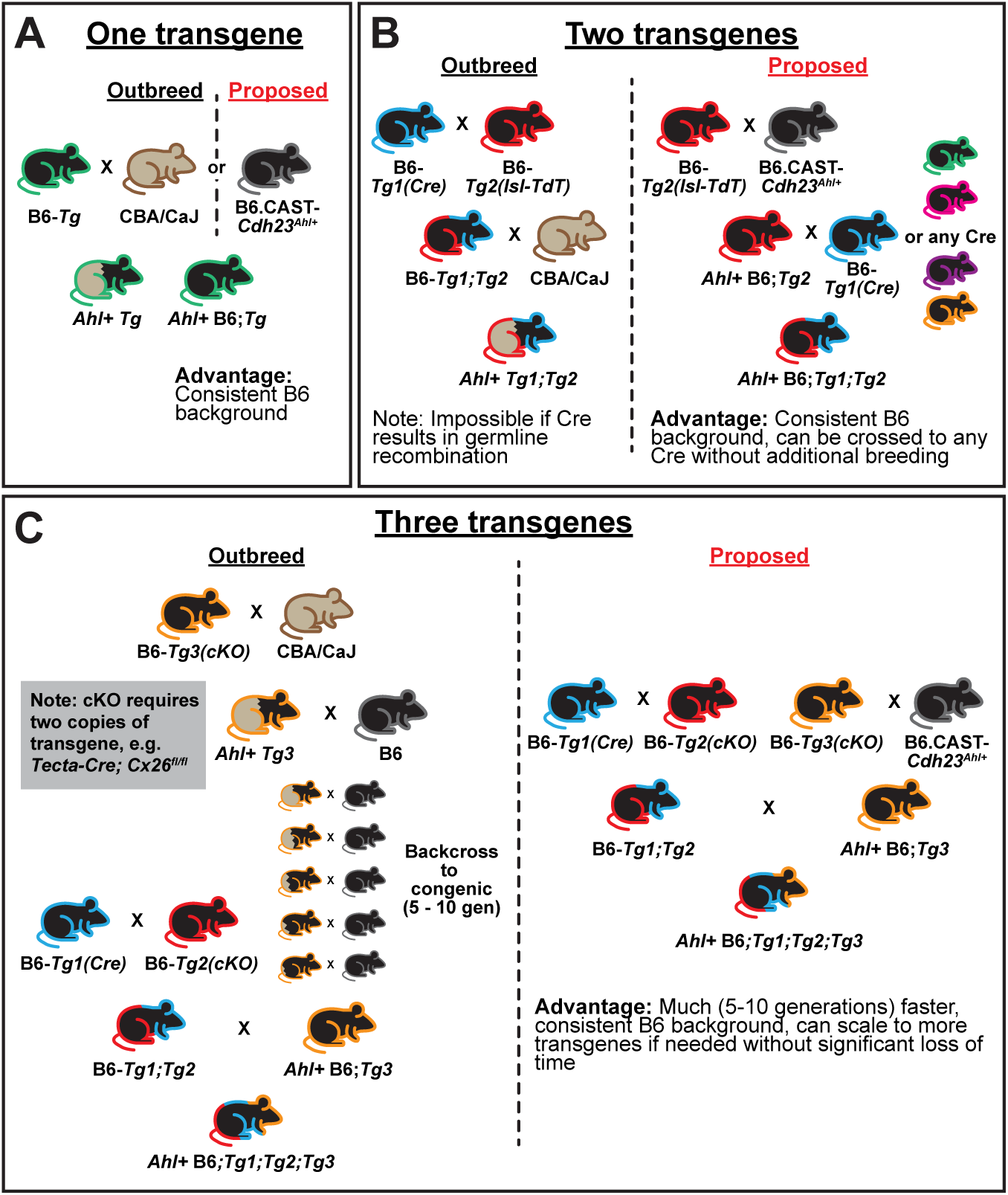
Breeding strategies to preserve hearing in transgenic mice. **A,** Schematic of breeding strategy for a single transgene, e.g. *Thy1-GCaMP6s* used in this study. Mouse fill indicates its strain; black is C57BL/6, brown is CBA/CaJ, brown/black indicates mixed strain. Colored outlines indicate the transgene. **B,** Schematic of breeding strategy for two transgenes, e.g. *Cre* and Cre-dependent (lox-stop-lox; lsl) reporter. Placing the reporter transgene on *Ahl+* background allows easy crossing to multiple Cre lines. **C,** Schematic of breeding strategy for three transgenes, e.g. *Cre* and conditional knockout (cKO) alleles (floxed/floxed).

### *C57BL/6* mice exhibit progressive hearing loss

Macroscopic widefield and two-photon imaging of *Thy1-GCaMP6s* mice expressing pan-neuronal GCaMP revealed an absence of low-threshold, cortical responses to high frequency sounds when mice were generated on a C57BL/6 background (Figure 1, 3). Previous studies have indicated that C57BL/6 mice contain an allelic variant in *Cdh23* that is responsible for the degradation of auditory responses and loss of hair cells, which is prevented by introducing a *Cdh23^c.753A^*^>^*^G^* single nucleotide substitution (Johnson et al., 2017) or by outbreeding and selecting for the *Ahl+* locus (Kane et al., 2012). Consistent with these observations, we observed robust low threshold responses at high frequencies (32 and 64kHz) by crossing *Thy1-GCaMP6s* mice to either *Ahl+* B6 mice (B6.CAST-Cdh23^Ahl+^/Kjn) or *Ahl+* CBA/CaJ mice. While responses to high frequencies were diminished on a pure C57BL/6 background, they were not absent as indicated by responses to tones presented at elevated sound levels (Figure 1E) and non-zero proportions of individual neurons exhibiting characteristic frequencies in that range (Figure 3H). These data are consistent with single-unit recordings in the primary auditory cortex of breeding age (8-12 weeks old) C57BL/6 females, which respond to pup calls that contain frequency components exclusively in the ultrasonic range (Marlin et al., 2015).

Two-photon imaging of cortical neurons in 2.5-month-old C57BL/6 mice revealed significant reorganization of neuronal tuning, with most neurons selectively responding to tones ≤ 16 kHz (Figure 4C, D). These data align with tonotopic map plasticity observed in cortical neurons, as measured with single-unit recordings in 3-month-old C57BL/6 mice (Willott et al., 1993). Similarly, ABRs recorded from 3-week-old C57BL/6 mice display elevated thresholds at high frequencies compared to CBA/CaJ controls (Zhang et al., 2013), with other studies indicating hearing deficits as early as 3 months of age (Kane et al., 2012; Ouagazzal et al., 2006; Zheng et al., 1999). These data strongly suggest that substantial hearing-loss has already occurred by early adulthood, prompting plasticity of the remaining sensory input. Additionally, cortical responses to tones decline sharply over the next four months in C57BL/6 mice (Figure 6A-C), which may prompt further reorganization due to reduced auditory drive (Willott et al., 1993). The progressive nature of this hearing loss inherently means that each mouse experiences a varying degree of hearing impairment during the first few months of life, a critical period for many experiments, raising concerns about its role as a potential confounding factor in many other studies.

### Potential confounds for systems neuroscience experiments

The use of C57BL/6 mice for auditory physiology and behavioral tasks has clear limitations. If the mice cannot hear or have an altered perception of the sound as part of the task, low performance may be mistakenly attributed to cognitive or motivational deficits, rather than a sensory deficit. For example, auditory cues are often paired with rewards in behavioral experiments (Guo et al., 2019; Olsen & Winder, 2009), and hearing loss can disrupt this association. If two tones are used, e.g. one low (16 kHz) and one high (32 kHz), behavioral differences at these two frequencies may reflect hearing loss, ultimately leading to incorrect assessments of reward-related learning without proper controls.

Progressive hearing loss can also impact behavioral experiments that use auditory cues to trigger a behavior (Breton-Provencher et al., 2022; Inagaki et al., 2018; Li et al., 2021; Robert et al., 2021). While the loss of peripheral input is a concern, this deficit could result in large-scale cross-modal plasticity (see Lee & Whitt, 2015 for review), with a decreased peripheral drive leading to compensatory changes across sensory systems (Petrus et al., 2014). Indeed, humans who experience deafness early in life display higher visual attention and processing, correlated with higher visually-related activity within the anatomical auditory cortex (Dye & Bavelier, 2013). Likewise, hearing deficits could introduce compensatory behaviors, such as a reliance on or adaptation to other sensory cues, that will ultimately disrupt the interpretation of those studies.

Pup-rearing behavior may also be impacted by hearing loss. During isolation or distress, newborn mouse pups signal their mothers with ultrasonic calls (>40 kHz). In C57BL/6 mice, first-time mothers exhibited more reliable and temporally precise action potential firing in response to these calls compared to virgin females (Marlin et al., 2015). While these findings might seem inconsistent with this study, pup distress calls can reach 80-90 dB SPL (Ehret, 2013), a sound level sufficient to elicit neural responses in *ahl* B6 mice (Figure 1E). However, variability in the prior study may reflect different degrees of hearing loss in the maternal population. Additionally, C57BL/6 mothers take longer to retrieve pups than CBA/CaJ mothers (Stevenson et al., 2021) and do not retrieve vocalizing pups at higher rates than non-vocalizing pups (Winters et al., 2023), indicating that other cues, such as pup odor (Okabe et al., 2013; Vinograd et al., 2017) or lower frequency calls (Ehret, 2006), may play a larger role in pup retrieval behaviors in C57BL/6 mice.

Environmental awareness is also largely driven by hearing, and loss of this sensation has been correlated with an increase in anxiety and stress levels in rodents (Lauer et al., 2018), potentially influencing behavioral responses in a range of experimental paradigms. Moreover, in humans hearing deficits are linked to social isolation, depression, and dementia (Blazer & Tucci, 2019; Garnefski & Kraaij, 2012; Mick et al., 2014), and behavioral correlates may exist in mice. Together, these studies indicate that the potential for hearing loss as a confounding factor impacts systems neuroscience as a whole, not just the auditory field. Therefore, researchers must carefully consider their choice of animal model and the potential impact of sensory deficits on their experimental outcomes to avoid misleading interpretations.

### Practical considerations for experiments

When using congenic *Ahl+* B6 mice to generate transgenic mouse models, we offer several suggestions. With a single transgene, the transgenic mouse can simply be crossed to B6.CAST-*Cdh23*^Ahl+^/Kjn rather than outbreeding to CBA/CaJ to maintain a consistent genetic background (Figure 7A). In most cases, we recommend breeding the mouse of interest to *Ahl+* homozygosity (*Cdh23*^Ahl/Ahl^). One allele is sufficient to protect against age-related hearing loss (Mianné et al., 2016; Perrin et al., 2013), so all offspring from these breeders will retain low threshold hearing. Maintaining breeders with *Cdh23*^Ahl/Ahl^ also limits extensive genotyping, as the offspring genotypes are predetermined. In experiments requiring Cre and Cre-dependent reporters, placing *Ahl+* alleles on the reporter line is usually advantageous, because those reporters can be used with multiple Cre lines (Figure 7B), as opposed to separately breeding those alleles onto each Cre line of interest. However, in cases of BAC transgenic Cre lines (which are generated with random insertion of Cre mediated by a *bac*terial artificial chromosome), it could be faster to reach homozygosity (3 total alleles) than in reporter lines (4 alleles, assuming homozygosity in *Cdh23* and reporter). When conditional deletion is required, our only suggestion is to reach *Ahl+* homozygosity on whichever mouse line has fewer total alleles to minimize the amount of total breeding required to generate the desired genotype (Figure 7C).

While progressive hearing loss is a major factor in implementing this strategy, we also expect less variation in behavior, as previous studies have demonstrated remarkable differences among mouse strains in common behavioral tasks (Brooks et al., 2005; Kim et al., 2017) and brain plasticity (Ranson et al., 2013). In this study, we observed that *Ahl+* B6.CBA mice exhibited lower signal correlations and higher proportions of tone-offset responding neurons than either *ahl* or *Ahl+* B6 mice (Supp Figure 3), indicating that these circuits operate differently than in C57BL/6 mice (Bowen et al., 2020). Ultimately, these differences would make comparisons between experiments using different strains extremely difficult.

We acknowledge that this approach requires startup costs related to obtaining, breeding, and ongoing maintenance of these mouse lines. However, the benefit of reduced variability in animal hearing will improve the interpretability of any study design by minimizing the risk of hearing loss as a confounding factor. In cases where this strategy is not feasible and C57BL/6 mice must be used, we strongly recommend using conventional hearing assessments (ABR) to include as an explanatory variable in any statistical models comparing groups, as *in vivo* contexts often rely on small sample sizes (∼5-6 mice/group) that can be significantly impacted by individual variability.

In summary, our work presents a streamlined strategy to mitigate progressive hearing loss in transgenic C57BL/6 mice. By introducing the *Ahl+* allelic variant through a cross with commercially-available congenic B6.CAST-Cdh23^Ahl+^/Kjn mice, we observed limited progressive hearing loss, as evidenced by their sustained low-threshold responses to high-frequency sounds up to 6 months of age. These approaches are easy to implement, scalable to more complex genetic models, and eliminate the need for labor-intensive genotyping techniques, thereby improving the reliability and interpretability of research across multiple neuroscience disciplines.

## ACKNOWLEDGMENTS

We thank Dr. Amit Agarwal for his helpful discussions about the *Cdh23* genotyping strategy, Lillian Choi for technical assistance, and members of the Kanold laboratory for discussions and comments on the manuscript. Funding was provided by grants from the NIH (T.A.B. – NIH F32DC019842, P.O.K. - NIH RO1DC009607).

## AUTHOR CONTRIBUTIONS

Conceptualization, T.A.B. and P.O.K.; Methodology, T.A.B.; Software: T.A.B., N.D., S.S.D.; Validation: T.A.B.; Formal Analysis, T.A.B., N.D, S.S.D.; Investigation, T.A.B., N.D., S.S.D., C.D.S.; Resources: T.A.B. and P.O.K.; Data Curation: T.A.B.; Writing – Original Draft, T.A.B., N.D.; Writing – Review & Editing, T.A.B., N.D., S.S.D., P.O.K.; Visualization: T.A.B.; Supervision: T.A.B. and P.O.K.; Project Administration: T.A.B. and P.O.K.; Funding Acquisition, T.A.B. and P.O.K.

## STAR METHODS

### RESOURCE AVAILABILITY

#### Lead Contact

Further information and requests for resources should be directed to and will be fulfilled by the lead contact, Patrick Kanold.

#### Materials availability

This study did not generate new unique reagents.

#### Data and code availability

Files generated following suite2p pre-processing (fluorescence, classification, neuropil fluorescence, etc.) and source code for analysis and figure generation will be available upon acceptance at the JHU Research Data Repository. DOIs will be listed in the key resources table. Any additional information required to reanalyze the data reported in this paper is available from the lead contact upon request.

### EXPERIMENTAL MODEL AND STUDY PARTICIPANT DETAILS

Both male and female mice aged between 2 and 6 months (as indicated for each experiment) were used for experiments. For the initial widefield imaging session, *ahl* B6 mice were (mean ± SD) 71 ± 17 days old, *Ahl+ B6* mice were 72 ± 18 days old, and *Ahl+* B6.CBA mice were 88 ± 9 days old. For 6-month-old widefield imaging session, *ahl* B6 mice were (mean ± SD) 191 ± 13 days old, *Ahl+* B6 mice were 196 ± 13 days old, and *Ahl+* B6.CBA mice were 188 ± 6 days old. For 2P imaging sessions, *ahl* B6 mice were (mean ± SD) 83 ± 17 days old, *Ahl+* B6 mice were 85 ± 24 days old, and *Ahl+* B6.CBA mice were 109 ± 10 days old.

We include sex as a factor for most analyses, which are reported if relevant and included in the supplemental statistics file for completeness. All animals were healthy (not-immunodeficient) and were only used for experiments detailed in this study. *Thy1-GCaMP6s* (GP4.3; Jax#: 024275) were crossed to *Cdh23*^Ahl/ahl^ heterozygotes (F1 offspring of B6.CAST-*Cdh23^Ahl+^*/Kjn (Jax#: 002756) and C57BL/6J (Jax#: 000664)) to generate *Ahl+* and control littermates. For *Ahl+* B6.CBA mice, F1 offspring of *Thy1-GCaMP6s* and *CBA/CaJ* mice were used.

Mice were group housed on a 12-hour light/dark cycle and were provided food ad libitum. This study was performed in accordance with the recommendations provided in the Guide for the Care and Use of Laboratory Animals of the National Institutes of Health. All experiments and procedures were approved by the Johns Hopkins Institutional Care and Use Committee. All surgery was performed under isoflurane anesthesia and every effort was made to minimize suffering.

### METHOD DETAILS

#### Genotyping *Cdh23* for *Ahl+* and *ahl* allelic variants

A KAPA2G HotStart kit (KK7352, Roche) was used for all genotyping. Briefly, DNA was extracted from tail clippings and amplified with PCR with the following primers: GTGCTGTTGGGCCTCCTTGC and GGGGTGGACCATGATCTATTTTGT. A 10ul reaction volume was used, comprised of 5 ul 2X KAPA2G mix, 2 ul of primer mix (10 uM final volume of each primer), 2 ul of DNAase, RNAase-free H_2_O, and 1 ul of extracted DNA. HotStart was initiated with 2 minutes at 95°C, followed by 32 repeats of 95°C-63°C-72°C steps (denaturing-annealing-extension; 10s each). A 2-minute final extension at 72°C was followed by a 4°C hold. Finally, 0.5 ul (500 units) of BsrI was added to the reaction, incubated at 65°C for 15 minutes, and run in 1.8% agarose gel for ∼15 minutes at 150V. Expected sizes are 284 and 664 base pairs for *ahl* and 948 bp for *Ahl+*.

#### Cranial window installation

##### Animal preparation

Before surgery, mice were injected with dexamethasone (2ug/g; delivered intramuscularly to the quadriceps of the hind paw) to limit brain swelling. Animals were anesthetized with isoflurane (Fluriso, VetOne) using a calibrated vaporizer. Anesthesia was induced at 4% isoflurane for 5 minutes and maintained at 1.5-2% for the duration of the surgery (∼ 1 hour). Body temperature was maintained at 36.5°C using an internal thermometer and feedback-controlled heating blanket (Harvard Apparatus 50-7212). Hair was removed from the scalp using a 5-minute application of Nair hair removal cream and thorough cleaning of the surgical area with three successive betadine and ethanol rinses. All surgical tools were sterilized before the first incision.

##### Cranial window and headpost implantation

Surgical scissors were used to remove the scalp from the dorsal aspect of the skull, starting slightly anterior to the ears to the Bregma suture, with lateral incisions approximately 3 mm to the right of and 5 mm to the left of the sagittal suture. The left temporalis muscle was retracted laterally to expose the squamosal suture and the posterior aspect of the jugal bone. Surgical calipers were used to lightly etch a 3.5 mm square encompassing the auditory cortex, with the posterior edge defined by the lamboid suture and the lateral edge defined by the squamosal suture. The “fold” of the parietal bone, running in the anterior to posterior direction was located roughly in the middle of this square. A 1 mm dental drill bit was used to bore a circular shape, followed by more delicate removal using a 0.5 mm bit. The skull was carefully removed and a modified coverslip consisting of a 4 mm circular cover glass adhered to a 3mm circular coverslip with optical glue was adhered to the skull initially with cyanoacrylate glue and permanently secured with dental cement, along with the headpost.

##### Post-operative care

Mice were kept on the surgical blanket until major muscle movements were observed, then transferred to a recovery cage placed on a circulating water blanket maintained at 36°C. Carprofen (Rimadyl; 5mg/kg IP) was administered immediately after the surgery and continued for the next 3 days. Dietgel (Clear H20) was provided in the cage during this recovery period.

#### Sound presentation and calibration

Half-second SAM tones (sinusoidal amplitude modulated, 10 Hz, cosine gated for first 10 ms) were delivered through a custom MATLAB script communicating with an RX6 Multifunction Processor (TDT). Tones were presented through an electrostatic speaker and driver (ES1 and ED1, TDT), with each tone individually calibrated to a 90 dB SPL reference with a Brüel and Kjær ultrasonic microphone.

#### Widefield imaging

Mice were head fixed in a holder under a custom widefield microscope. The field of view was illuminated with a 470 nm LED (M470L3, Thorlabs) through a 4x air objective (UplanSApo 4×/0.16, Olympus) mounted at ∼45° from vertical. The light path was separated via a long pass dichroic mirror (MD499-FITC, Thorlabs) to illuminate the sample, and the light was collected via a sCMOS camera (pco.edge 4.2, Excelitas Technologies) following emission filter (AT535/40m, Chroma) and tube lens focusing (AC508-150A, Thorlabs). The camera and sound stimuli were triggered externally (USB-6259, National Instruments) at 30 Hz. Images were digitized at 330 x 330 pixels that encompassed the entire cranial window (∼3mm^2^).

#### Widefield data processing and analysis

Suite2p (Pachitariu et al., 2017) was used to register (non-rigid, 128 x 128 block size) widefield images. Default parameters were used except for the sampling rate (fs), which was changed to 30 to match our acquisition parameters. Images were then downsampled from 330 pixels^2^ to 100 pixels^2^ to hasten further calculations. Raw fluorescence traces were then unmixed, forming a matrix of size X x Y x T x F x A x R, where X = width of the frame in pixels (100), Y = height of frame in pixels (100), T = time of the trial period (75 frames), F = number of frequencies (5), A = number of attenuations or sound levels (4), and R = repeats (10).

The baseline (F_o_) was calculated by averaging the 30 frames (1s) prior to tone onset for each trial. This baseline was used to normalize each pixel’s signal by calculating ΔF/F as (F - F_o_) / F_o._ For each tone and sound level response, we averaged the window between 10 and 15 frames after tone onset and used this as the sound response amplitude (visualized in images Figure 1D,G). We then averaged only pixels with ΔF/F in the 99^th^ percentile (visualized as traces in Figure 1D,G). The threshold was calculated based on these responses, using a paired t-test between the baseline and response amplitude, with alpha = 0.05 and no correction for multiple comparisons. Thresholds were defined as the sound with the first statistically significant response, or 110 dB SPL if there was no detectable response at the highest sound level presented.

#### Two-photon imaging

Mice were head-fixed in a holder under an Ultima 2Pplus microscope (Bruker). This microscope features a rotatable objective to achieve imaging of the auditory cortex with animals in a normal stationary position. A 920 nm excitation laser (Chameleon Discovery NX, Coherent) was used in conjunction with a 16x objective (CFI75 LWD 16X W, Nikon) and resonant-galvo scanning to capture images at ∼15 Hz at 1024 pixel^2^ resolution, resulting in a field of view of ∼1.1 mm^2^ that was targeted to primary auditory cortex with widefield imaging maps (Figure 1C). The frame out signal from the microscope was used to trigger sound stimuli.

#### Two-photon data processing and analysis

##### Registration, fluorescence extraction, and DF/F calculation

Suite2p (Pachitariu et al., 2017) was used to register (non-rigid, 128 x 128 block size) and extract fluorescence from regions of interest (ROIs) and surrounding neuropil. Default parameters were used except for the sampling rate (fs), which was changed to 15 to match our acquisition parameters.

For each identified ROI, the raw fluorescence signal over time,*F_ROI_*, was extracted. The fluorescence signal used for analysis,*F*, was calculated by performing neuropil (NP) subtraction on the raw fluorescence signals of each ROI: *F* = *F_ROI_* − (α × *F_NP_*) where α = 0.7, to decrease the contamination from neuropil fluorescence. To correct for slow drifts that occur over the length of the imaging session, we calculated the baseline (F_o_) using a moving-window approach. The baseline was defined as the 10^th^ percentile of fluorescence values within a window of 500 frames with 50 frame steps. The resulting baseline was linearly interpolated to match the size of the original input. Finally, this baseline was used to normalize the neuronal signal by calculating ΔF/F as (F - F_o_) / F_o_.

##### Determination of sound-responsive neurons

To determine if a neuron was sound-responsive, we created a linear model to explain the signal amplitude using a response variable that encoded the signal phase (baseline, onset, or offset), the frequency, the attenuation, and the interactions between these variables (signal_amp ∼ C(response_variable) + C(frequency) + C(attenuation) + C(frequency):C(attenuation):C(response_variable)). The response amplitude was calculated as the mean response during the 1 second before the sound was played (baseline), between frames 5 and 9 after tone onset (onset, approximately 330-600 ms after tone onset), and between frames 5 and 9 after tone offset (offset). We considered the neuron sound-responsive if the response variable explained the variance at an α < 0.01, following criteria used in previous studies (Bowen et al., 2020; Winkowski & Kanold, 2013). If the response variable was significant at this level, we performed post-hoc t-tests with Benjamini/Hochberg correction to compare the means of each response phase. If the onset amplitude was significantly higher than the baseline amplitude, the neurons were classified as “Onset” neurons. If the offset amplitude was significantly higher than the onset amplitude, the neurons were classified as “Offset” neurons. If both properties were present, the neuron was classified as an “Onset/Offset” neuron. For calculating the proportion of sound offset-responsive neurons, both “Offset” and “Onset/Offset” neurons were grouped as “Offset” neurons.

##### Best frequency, characteristic frequency, and bandwidth calculation

For all calculations described in this section, only the onset response amplitudes for “Onset” and “Onset/Offset” neurons were used. For every frequency and attenuation level, we performed post-hoc t-tests with multiple comparisons controlled via Benjamini-Hochberg procedure with a false discovery rate (α = 0.2). Only frequency and sound levels that showed a statistically significant response were included for subsequent analysis.

Bandwidth was defined as log_2_(f_max_/f_min_), with f_max_ and f_min_ representing the maximum and minimum frequency with a significant response. Neurons with non-continuous frequency responses (i.e. multiple peaks) were excluded from the analysis. Best frequency was defined as the frequency with the highest amplitude response, regardless of sound level. Characteristic frequency was defined as the frequency with the highest amplitude response at the lowest sound level.

##### Signal and Noise Correlations

Signal and noise correlations were calculated following methods from previous studies (Bowen et al., 2020; Winkowski & Kanold, 2013). To measure signal correlations, we first averaged ΔF/F responses across all repeats for each frequency and sound level. We calculated the correlation coefficient over a 2-second window (30 frames) after tone onset for each frequency and sound level between all pairs of sound-responsive neurons. These correlation values were then averaged to obtain a signal correlation value for each neuron pair. Finally, we averaged the correlation values across all pairs to get the overall signal correlation for each animal.

To measure noise correlations, which assess the correlations between neurons that are independent of their response to the sound, we first averaged ΔF/F responses across all repeats for each frequency and sound level and then subtracted this from each repeat. For each neuron pair, we computed the correlation coefficient on a repeat-by-repeat basis over a 1-second window (30 frames) after tone onset. Finally, we averaged these correlation coefficients across all neuron pairs, frequency levels, sound levels, and repeats to obtain the overall noise correlation for each animal.

##### Linear Discriminant Analysis

ΔF/F traces were unmixed, forming a matrix of size N x T x F x A x R, where N = number of neurons, T = time of the trial period (75 frames), F = number of frequencies (5), A = number of attenuations or sound levels (4), and R = repeats (10). For each trial, we calculated the average response over a 1-second (15 frames) following tone-onset, resulting in an N x 1 x F x A x R array that was then reshaped into an N x (F x A x R) array.

The (F x A x R) dataset was randomly split into 90% training and 10% test data. For each split, Principal Component Analysis was fit on the training subset, and linear features explaining at least 90% of the data variance extracted with this fit were used to train a linear discriminant analysis (LDA) classifier to classify neural trials by the corresponding tone and sound levels. The PCA features and LDA classifier were deployed on the testing subset to evaluate classification performance. This procedure was repeated 10 times to cover the entire dataset, and the average accuracy was reported. Full implementation details are available in the provided code.

### QUANTIFICATION AND STATISTICAL ANALYSIS

All statistics and corrections for multiple comparisons were performed in Python with the statsmodels (for linear models) and scipy (for post-hoc testing) packages. All statistical details, including the exact value of n, what n represents, and which statistical test was performed, can be found in the figure legends and/or within the figure panels. Full statistical details for every figure are included in the supplement. Data in plots are presented as mean ± standard error of the mean, unless indicated otherwise. For single comparisons, significance was defined as *p* ≤ 0.05. When multiple comparisons were made, the Benjamini-Hochberg correction was used to adjust p-values accordingly to lower the probability of type I errors. For multiple condition datasets, an N-way ANOVA analysis was performed, and, if relevant variables were statistically significant, followed by t-test comparison with Benjamini-Hochberg correction.

## FIGURE LEGENDS

**Supplementary Figure 1.**
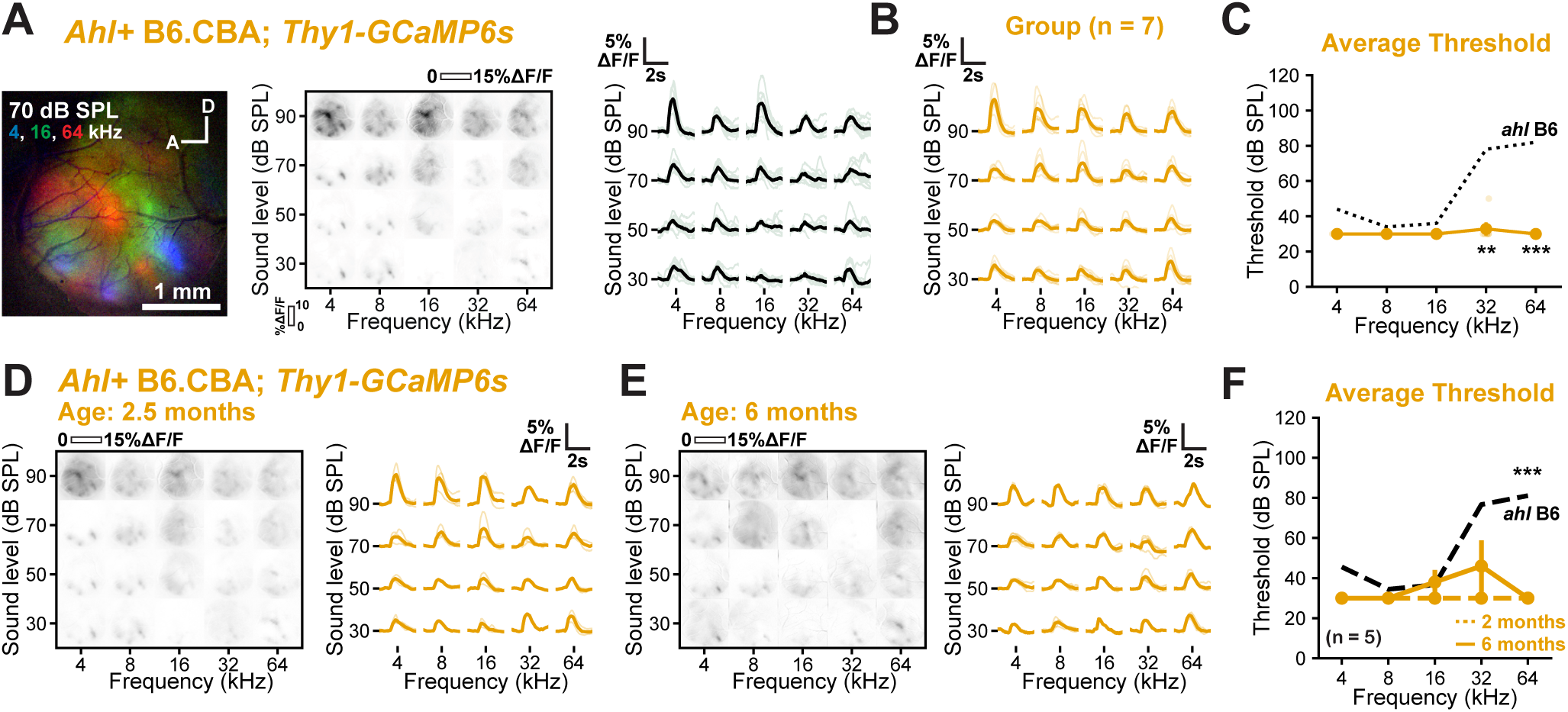
Widefield imaging in *Ahl+* B6.CBA mice reveals low threshold responses to a broad range of frequencies. **A,** (left) Merged fluorescence responses to 4 (blue), 16 (green), and 64kHz (red) at 70 dB SPL in P70 *Ahl+* B6.CBA; *Thy1-GCaMP6s* mice. (middle) Plot of fluorescence changes over the imaging field to varying frequency (x-axis) and sound levels (y-axis). (right) Plot of fluorescence during sound presentation from an individual mouse. Grey traces are individual trials, black traces are the average. **B,** Plot of fluorescence during sound presentation from all *Ahl+* B6.CBA; *Thy1-GCaMP6s* mice. Light traces are individual mice, dark traces are the group average, n = 7 mice. **C,** Plot of average fluorescence detection threshold for each frequency presented, n = 7 mice. Three-way ANOVA (frequency: *F(4,110)* = 16.2*, p* = 1.8e-10; sex: *F(1,110)* = 4.76*, p* = 3.1e-02; genotype: *F(2,110)* = 61.7*, p* = 1.1e-18; interaction: *F(22,110)* = 5.13*, p* = 3.2e-09) followed by planned comparisons with t-tests controlled with Benjamini-Hochberg FDR. **: *p* < 0.01, ***: *p* < 0.001. **D,** (left) Plot of fluorescence changes in 2.5-month-old *Thy1-GCaMP6s* (*Ahl+* B6.CBA) mice over the imaging field to varying frequency (x-axis) and sound levels (y-axis). (right) Plot of fluorescence during sound presentation across animals; grey traces are individual mice, black traces are the average, n = 5 mice. **E,** (left) Plot of fluorescence changes in 6-month-old *Thy1-GCaMP6s* (*Ahl+* B6.CBA) mice over the imaging field to varying frequency (x-axis) and sound levels (y-axis). (right) Plot of fluorescence during sound presentation across animals; grey traces are individual mice, black traces are the average, n = 5 mice. **F,** Plot of average threshold as a function of frequency and time point. Dashed lines indicate measurements at 2.5 months, solid lines indicate measurements at 6 months, n = 5 mice. Three-way ANOVA (frequency: *F(4,30)*=4.80*, p* = 4.1e-3; timepoint: *F(1,30)*=10.8*, p* = 2.6e-3; sex: *F(1,30)*=16.2*, p* = 3.6e-4, interaction: *F(13,30)*=7.15*, p* = 5.0e-6) with post-hoc paired t-tests with Benjamini-Hochberg FDR correction. n.s.: not significant.

**Supplementary Figure 2.**
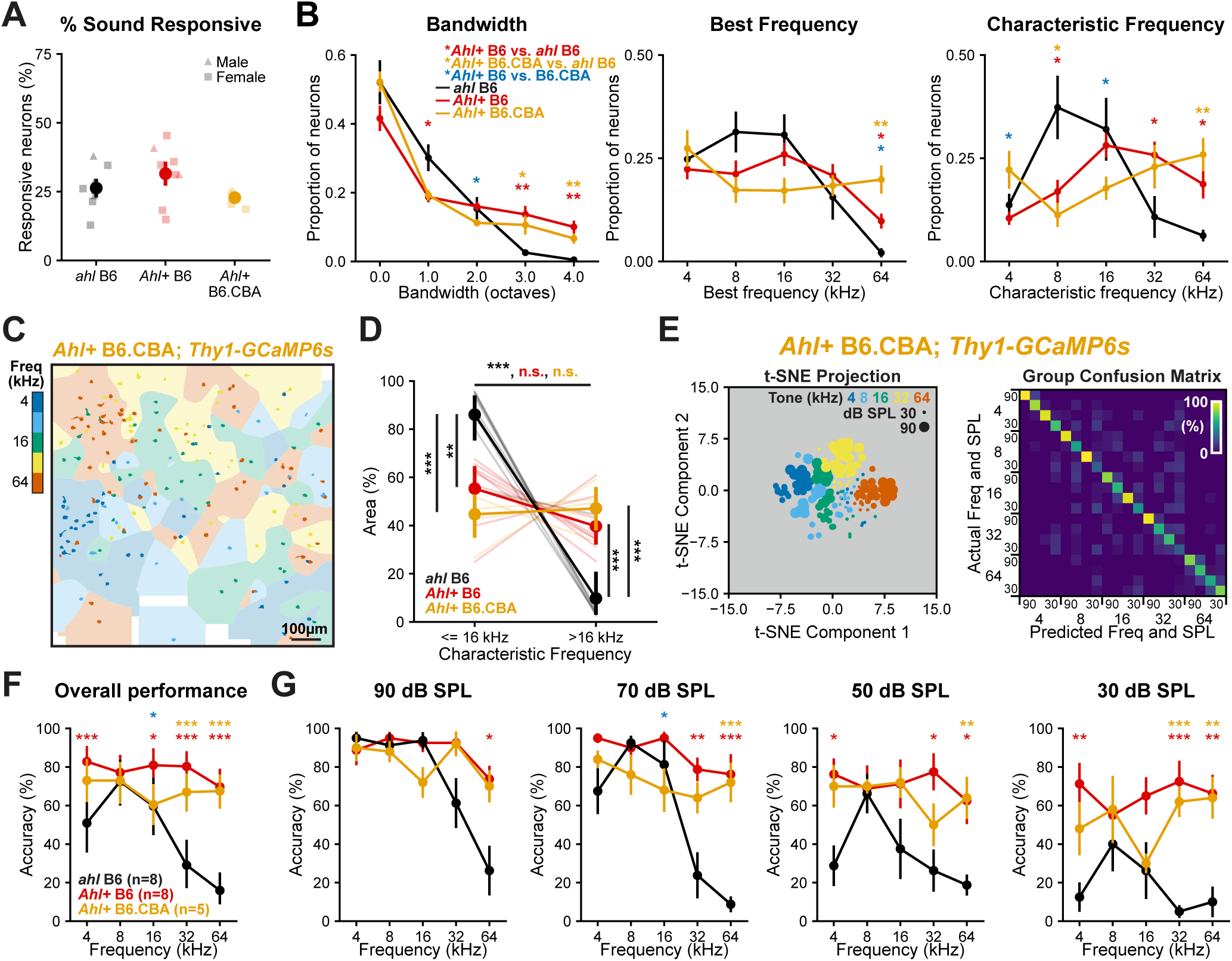
Neuronal responses in *Ahl+* B6.CBA are similar to *Ahl+* B6 mice. **A,** Plot of the proportion of sound-responsive neurons within each genotype. Light markers indicate individuals, dark markers are mean ± SEM, n = 8 mice for *ahl* and *Ahl+* B6 mice, n = 5 mice for *Ahl+* B6.CBA mice. Two-way ANOVA (genotype: *F(2,15) =* 2.42, *p*=0.12; sex: *F(1,15)* = 1.95*, p* = 0.18, interaction: *F(2,15)* = 0.00*, p* = 0.99). n.s.: not significant. **B,** Plots of the proportion of neurons as a function of bandwidth, best frequency, and characteristic frequencies across genotypes. Points are mean ± SEM, n = 8 mice for *ahl* and *Ahl+* B6 mice, n = 5 mice for *Ahl+* B6.CBA mice. Two-way ANOVAs with genotype and measured characteristic (see Supp Statistics for values), followed by post-hoc t-tests with Benjamini-Hochberg FDR correction. *: *p* < 0.05, **: *p* < 0.01, ***: *p* < 0.001. Comparisons between *Ahl+* B6 and *ahl* B6 are indicated with red stars, *Ahl+* B6.CBA and *ahl* B6 with yellow stars, and *Ahl+* B6 and *Ahl+* B6.CBA with blue stars. **C,** Exemplar characteristic frequency maps and assigned areas for *Ahl+* B6.CBA mice. White areas on the map indicate regions where no characteristic frequency was assigned due to a lack of neurons within the area. **D,** Plot of normalized area as a function of characteristic frequency and genotype. Light lines are individual animals, dark lines are mean ± SEM, n = 8 mice for *ahl* and *Ahl+* B6, *n* = 5 for *Ahl+* B6.CBA. Two-way ANOVA (characteristic frequency: *F(1,36)* = 72.98*, p* < 0.001; genotype: *F(2,36) = 0.08, p* = 0.92; interaction: *F(2,36)* = 34.6, *p* < 0.001), followed by post-hoc t-tests with Benjamini-Hochberg FDR correction. n.s: not significant, **: *p* < 0.01, ***: *p* < 0.001. **E,** (left) Low-dimensional representation (t-SNE) of neuronal population response from an individual *Ahl+* B6.CBA mouse with each marker representing a single trial. (right) Confusion matrix of classifier performance from a single animal. **F,** Plot of overall classifier performance as a function of frequency, n = 8 mice for *ahl* and *Ahl+* B6 mice, n = 5 mice for *Ahl+* B6.CBA mice. Three-way ANOVA (genotype: *F(2,360)* = 76.7*, p* < 0.001; frequency: *F(4,360)* = 15.2*, p* < 0.001; sound level: *F(3,360)* = 47.8*, p* < 0.001; interaction: *F(50,360)*=2.89*, p* < 0.001), all attenuation levels included for each frequency in post-hoc t-tests with Benjamini-Hochberg FDR correction. *: *p* < 0.05, **: *p* < 0.01, ***: *p* < 0.001. Comparisons between *Ahl+* B6 and *ahl* B6 are indicated with red stars, *Ahl+* B6.CBA and *ahl* B6 with yellow stars, and *Ahl+* B6 and *Ahl+* B6.CBA with blue stars. **G,** Plot of classifier performance as a function of frequency (x-axis) and sound level (plots arranged from highest sound level to lowest sound level). n = 8 mice for *ahl* and *Ahl+* B6 mice, n = 5 mice for *Ahl+* B6.CBA mice. The same three-way ANOVA is reported in **F,** post-hoc t-tests with Benjamini-Hochberg FDR correction. *: *p* < 0.05, **: *p* < 0.01, ***: *p* < 0.001. Comparisons between *Ahl+* B6 and *ahl* B6 are indicated with red stars, *Ahl+* B6.CBA and *ahl* B6 with yellow stars, and *Ahl+* B6 and *Ahl+* B6.CBA with blue stars.

**Supplementary Figure 3.**
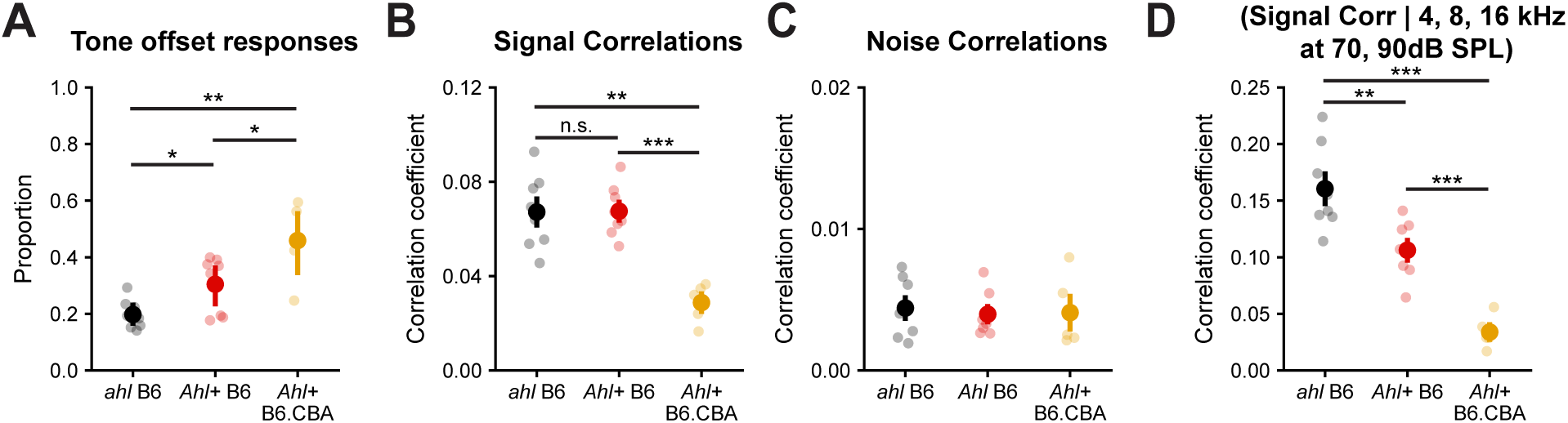
Network-level analysis of neuronal responses across genotypes. **A,** Plot of the proportion of tone offset responses as a function of genotype, *n* = 8 mice for *ahl* and *Ahl+* B6, *n* = 5 for *Ahl+* B6.CBA. Two-way ANOVA (genotype: *F(2,15)* = 11.3, *p* = 0.001; sex: *F(1,15)* = 0.02, *p* = 0.89; interaction: *F(2,15)* = 1.18, *p* = 0.33), followed by post-hoc t-tests with Benjamini-Hochberg FDR correction. *: *p* < 0.05, **: *p* < 0.01 **B,** Plot of signal correlations among sound-responsive neurons as a function of genotype, *n* = 8 mice for *ahl* and *Ahl+* B6, *n* = 5 for *Ahl+* B6.CBA. Two-way ANOVA (genotype: *F(2,15)* = 16.7, *p* < 0.001; sex: *F(1,15)* = 0.42, *p* = 0.52; interaction: *F(2,15)* = 0.50, *p* = 0.61), followed by post-hoc t-tests with Benjamini-Hochberg FDR correction. **: *p* < 0.01, ***: *p* < 0.001. **C,** Plot of noise correlations among sound-responsive neurons as a function of genotype, *n* = 8 mice for *ahl* and *Ahl+* B6, *n* = 5 for *Ahl+* B6.CBA. Two-way ANOVA (genotype: *F(2,15)* = 0.06, *p* = 0.93; sex: *F(1,15)* = 1.11, *p* = 0.31; interaction: *F(2,15)* = 0.97, *p* = 0.40). **D,** Plot of signal correlations among sound-responsive neurons as a function of genotype, conditioned on low frequencies (4, 8, 16 kHz) and higher sound levels (70 and 90 dB SPL), *n* = 8 mice for *ahl* and *Ahl+* B6, *n*= 5 for *Ahl+* B6.CBA. Two-way ANOVA (genotype: *F(2,15)* = 41.0, *p* < 0.001; sex: *F(1,15)* = 3.28, *p* < 0.001; interaction: *F(2,15)* = 2.95, *p* = 0.08), followed by post-hoc t-tests with Benjamini-Hochberg FDR correction. **: *p* < 0.01, ***: *p* < 0.001.

## REFERENCES

Blazer, D. G., & Tucci, D. L. (2019). Hearing loss and psychiatric disorders: A review. Psychological Medicine, 49(6), 891–897. 10.1017/S0033291718003409

Bowen, Z., Winkowski, D. E., & Kanold, P. O. (2020). Functional organization of mouse primary auditory cortex in adult C57BL/6 and F1 (CBAxC57) mice. Scientific Reports, 10(1), 10905. 10.1038/s41598-020-67819-4

Breton-Provencher, V., Drummond, G. T., Feng, J., Li, Y., & Sur, M. (2022). Spatiotemporal dynamics of noradrenaline during learned behaviour. Nature, 606(7915), 732–738. 10.1038/s41586-022-04782-2

Brooks, S. P., Pask, T., Jones, L., & Dunnett, S. B. (2005). Behavioural profiles of inbred mouse strains used as transgenic backgrounds. II: Cognitive tests. *Genes*, Brain and Behavior, 4(5), 307–317. 10.1111/j.1601-183X.2004.00109.x

Chambers, A. R., Resnik, J., Yuan, Y., Whitton, J. P., Edge, A. S., Liberman, M. C., & Polley, D. B. (2016). Central gain restores auditory processing following near-complete cochlear denervation. Neuron, 89(4), 867–879. 10.1016/j.neuron.2015.12.041

Dana, H., Chen, T.-W., Hu, A., Shields, B. C., Guo, C., Looger, L. L., Kim, D. S., & Svoboda, K. (2014). Thy1-GCaMP6 transgenic mice for neuronal population imaging in vivo. PLOS ONE, 9(9), e108697. 10.1371/journal.pone.0108697

de Vries, S. E. J., Lecoq, J. A., Buice, M. A., Groblewski, P. A., Ocker, G. K., Oliver, M., Feng, D., Cain, N., Ledochowitsch, P., Millman, D., Roll, K., Garrett, M., Keenan, T., Kuan, L., Mihalas, S., Olsen, S., Thompson, C., Wakeman, W., Waters, J., … Koch, C. (2020). A large-scale standardized physiological survey reveals functional organization of the mouse visual cortex. Nature Neuroscience, 23(1), 138–151. 10.1038/s41593-019-0550-9

Dye, M. W. G., & Bavelier, D. (2013). Visual attention in deaf humans: A neuroplasticity perspective. In A. Kral, A. N. Popper, & R. R. Fay (Eds.), Deafness (pp. 237–263). Springer. 10.1007/2506_2013_9

Ehret, G. (2006). Common rules of communication sound perception. In Behaviour and neurodynamics for auditory communication (pp. 85–114).

Ehret, G. (2013). Sound communication in house mice: Emotions in their voices and ears? In E. Altenmüller, S. Schmidt, & E. Zimmermann (Eds.), Evolution of emotional communication: From sounds in nonhuman mammals to speech and music in man (pp. 63–74). Oxford University Press. 10.1093/acprof:oso/9780199583560.003.0004

Frisina, R. D., Singh, A., Bak, M., Bozorg, S., Seth, R., & Zhu, X. (2011). F1 (CBA × C57) mice show superior hearing in old age relative to their parental strains: Hybrid vigor or a new animal model for “Golden Ears”? Neurobiology of Aging, 32(9), 1716–1724. 10.1016/j.neurobiolaging.2009.09.009

Garnefski, N., & Kraaij, V. (2012). Cognitive coping and goal adjustment are associated with symptoms of depression and anxiety in people with acquired hearing loss. International Journal of Audiology, 51(7), 545–550. 10.3109/14992027.2012.675628

Guo, L., Weems, J. T., Walker, W. I., Levichev, A., & Jaramillo, S. (2019). Choice-selective neurons in the auditory cortex and in its striatal target encode reward expectation. The Journal of Neuroscience, 39(19), 3687–3697. 10.1523/JNEUROSCI.2585-18.2019

Inagaki, H. K., Inagaki, M., Romani, S., & Svoboda, K. (2018). Low-dimensional and monotonic preparatory activity in mouse anterior lateral motor cortex. The Journal of Neuroscience, 38(17), 4163–4185. 10.1523/JNEUROSCI.3152-17.2018

International Mouse Knockout Consortium. (2007). A mouse for all reasons. Cell, 128(1), 9–13. 10.1016/j.cell.2006.12.018

Ison, J., & Allen, P. (2004). Low-frequency tone pips elicit exaggerated startle reflexes in C57BL/6J mice with hearing loss. Journal of the Association for Research in Otolaryngology, 4, 495–504. 10.1007/s10162-002-3046-2

Johnson, K. R., Tian, C., Gagnon, L. H., Jiang, H., Ding, D., & Salvi, R. (2017). Effects of Cdh23 single nucleotide substitutions on age-related hearing loss in C57BL/6 and 129S1/Sv mice and comparisons with congenic strains. Scientific Reports, 7(1), 44450. 10.1038/srep44450

Kane, K. L., Longo-Guess, C. M., Gagnon, L. H., Ding, D., Salvi, R. J., & Johnson, K. R. (2012). Genetic background effects on age-related hearing loss associated with Cdh23 variants in mice. Hearing Research, 283(1–2), 80–88. 10.1016/j.heares.2011.11.007

Keithley, E. M., Canto, C., Zheng, Q. Y., Fischel-Ghodsian, N., & Johnson, K. R. (2004). Age-related hearing loss and the ahl locus in mice. Hearing Research, 188(1–2), 21–28. 10.1016/S0378-5955(03)00365-4

Kim, J. W., Nam, S. M., Yoo, D. Y., Jung, H. Y., Kim, I. Y., Hwang, I. K., Seong, J. K., & Yoon, Y. S. (2017). Comparison of adult hippocampal neurogenesis and susceptibility to treadmill exercise in nine mouse strains. Neural Plasticity, 2017(1), 5863258. 10.1155/2017/5863258

Lauer, A. M., Larkin, G., Jones, A., & May, B. J. (2018). Behavioral animal model of the emotional response to tinnitus and hearing loss. Journal of the Association for Research in Otolaryngology, 19(1), 67–81. 10.1007/s10162-017-0642-8

Lee, H.-K., & Whitt, J. L. (2015). Cross-modal synaptic plasticity in adult primary sensory cortices. Current Opinion in Neurobiology, 35, 119–126. 10.1016/j.conb.2015.08.002

Li, Z., Wei, J.-X., Zhang, G.-W., Huang, J. J., Zingg, B., Wang, X., Tao, H. W., & Zhang, L. I. (2021). Corticostriatal control of defense behavior in mice induced by auditory looming cues. Nature Communications, 12(1), 1040. 10.1038/s41467-021-21248-7

Liu, J., & Kanold, P. O. (2021). Diversity of receptive fields and sideband inhibition with complex thalamocortical and intracortical origin in L2/3 of mouse primary auditory cortex. The Journal of Neuroscience, 41(14), 3142–3162. 10.1523/JNEUROSCI.1732-20.2021

Liu, J., Whiteway, M. R., Sheikhattar, A., Butts, D. A., Babadi, B., & Kanold, P. O. (2019). Parallel processing of sound dynamics across mouse auditory cortex via spatially patterned thalamic inputs and distinct areal intracortical circuits. Cell Reports, 27(3), 872–885.e7. 10.1016/j.celrep.2019.03.069

Luo, L., Ambrozkiewicz, M. C., Benseler, F., Chen, C., Dumontier, E., Falkner, S., Furlanis, E., Gomez, A. M., Hoshina, N., Huang, W.-H., Hutchison, M. A., Itoh-Maruoka, Y., Lavery, L. A., Li, W., Maruo, T., Motohashi, J., Pai, E. L.-L., Pelkey, K. A., Pereira, A., … Craig, A. M. (2020). Optimizing nervous system-specific gene targeting with Cre driver lines: Prevalence of germline recombination and influencing factors. Neuron, 106(1), 37–65.e5. 10.1016/j.neuron.2020.01.008

Madisen, L., Zwingman, T. A., Sunkin, S. M., Oh, S. W., Zariwala, H. A., Gu, H., Ng, L. L., Palmiter, R. D., Hawrylycz, M. J., Jones, A. R., Lein, E. S., & Zeng, H. (2010). A robust and high-throughput Cre reporting and characterization system for the whole mouse brain. Nature Neuroscience, 13(1), 133–140. 10.1038/nn.2467

Marlin, B. J., Mitre, M., D’amour, J. A., Chao, M. V., & Froemke, R. C. (2015). Oxytocin enables maternal behaviour by balancing cortical inhibition. Nature, 520(7548), 499–504. 10.1038/nature14402

McGill, M., Hight, A. E., Watanabe, Y. L., Parthasarathy, A., Cai, D., Clayton, K., Hancock, K. E., Takesian, A., Kujawa, S. G., & Polley, D. B. (2022). Neural signatures of auditory hypersensitivity following acoustic trauma. eLife, 11, e80015. 10.7554/eLife.80015

Mianné, J., Chessum, L., Kumar, S., Aguilar, C., Codner, G., Hutchison, M., Parker, A., Mallon, A.-M., Wells, S., Simon, M. M., Teboul, L., Brown, S. D. M., & Bowl, M. R. (2016). Correction of the auditory phenotype in C57BL/6N mice via CRISPR/Cas9-mediated homology directed repair. Genome Medicine, 8(1), 16. 10.1186/s13073-016-0273-4

Mick, P., Kawachi, I., & Lin, F. R. (2014). The association between hearing loss and social isolation in older adults. Otolaryngology–Head and Neck Surgery, 150(3), 378–384. 10.1177/0194599813518021

Mikaelian, D. O., Warfield, D., & Norris, O. (1974). Genetic progressive hearing loss in the C57/M6 mouse: Relation of behaviorial responses to cochlear anatomy. Acta Oto-Laryngologica, 77(1–6), 327–334. 10.3109/00016487409124632

Noben-Trauth, K., Zheng, Q. Y., & Johnson, K. R. (2003). Association of cadherin 23 with polygenic inheritance and genetic modification of sensorineural hearing loss. Nature Genetics, 35(1), 21–23.

Noreña, A. J., Moffat, G., Blanc, J. L., Pezard, L., & Cazals, Y. (2010). Neural changes in the auditory cortex of awake guinea pigs after two tinnitus inducers: Salicylate and acoustic trauma. Neuroscience, 166(4), 1194–1209. 10.1016/j.neuroscience.2009.12.063

Okabe, S., Nagasawa, M., Kihara, T., Kato, M., Harada, T., Koshida, N., Mogi, K., & Kikusui, T. (2013). Pup odor and ultrasonic vocalizations synergistically stimulate maternal attention in mice. Behavioral Neuroscience, 127(3), 432–438. 10.1037/a0032395

Olsen, C. M., & Winder, D. G. (2009). Operant sensation seeking engages similar neural substrates to operant drug seeking in C57 mice. Neuropsychopharmacology, 34(7), 1685–1694. 10.1038/npp.2008.226

Ouagazzal, A.-M., Reiss, D., & Romand, R. (2006). Effects of age-related hearing loss on startle reflex and prepulse inhibition in mice on pure and mixed C57BL and 129 genetic background. Behavioural Brain Research, 172(2), 307–315. 10.1016/j.bbr.2006.05.018

Pachitariu, M., Stringer, C., Dipoppa, M., Schröder, S., Rossi, L. F., Dalgleish, H., Carandini, M., & Harris, K. D. (2017). Suite2p: Beyond 10,000 neurons with standard two-photon microscopy. bioRxiv. 10.1101/061507

Perrin, B. J., Strandjord, D. M., Narayanan, P., Henderson, D. M., Johnson, K. R., & Ervasti, J. M. (2013). β-actin and fascin-2 cooperate to maintain stereocilia length. The Journal of Neuroscience, 33(19), 8114–8121. 10.1523/JNEUROSCI.0238-13.2013

Petrus, E., Isaiah, A., Jones, A. P., Li, D., Wang, H., Lee, H.-K., & Kanold, P. O. (2014). Crossmodal induction of thalamocortical potentiation leads to enhanced information processing in the auditory cortex. Neuron, 81(3), 664–673. 10.1016/j.neuron.2013.11.023

Ranson, A., Sengpiel, F., & Fox, K. (2013). The role of GluA1 in ocular dominance plasticity in the mouse visual cortex. The Journal of Neuroscience, 33(38), 15220–15225. 10.1523/JNEUROSCI.2078-13.2013

Robert, B., Kimchi, E. Y., Watanabe, Y., Chakoma, T., Jing, M., Li, Y., & Polley, D. B. (2021). A functional topography within the cholinergic basal forebrain for encoding sensory cues and behavioral reinforcement outcomes. eLife, 10, e69514. 10.7554/eLife.69514

Romero, S., Hight, A. E., Clayton, K. K., Resnik, J., Williamson, R. S., Hancock, K. E., & Polley, D. B. (2020). Cellular and widefield imaging of sound frequency organization in primary and higher order fields of the mouse auditory cortex. *Cerebral Cortex (New York*, NY), 30(3), 1603–1622. 10.1093/cercor/bhz190

Sinclair, J. L., Barnes-Davies, M., Kopp-Scheinpflug, C., & Forsythe, I. D. (2017). Strain-specific differences in the development of neuronal excitability in the mouse. Hearing Research. 10.1016/j.heares.2017.08.004

Stevenson, P., Casenhiser, D. M., Lau, B. Y. B., & Krishnan, K. (2021). Systematic analysis of goal-related movement sequences during maternal behavior in a female mouse model for Rett syndrome. The European Journal of Neuroscience, 54(2), 4528–4549. 10.1111/ejn.15327

Sultana, R., Ogundele, O. M., & Lee, C. C. (2019). Contrasting characteristic behaviours among common laboratory mouse strains. Royal Society Open Science, 6(6), 190574. 10.1098/rsos.190574

Vinograd, A., Fuchs-Shlomai, Y., Stern, M., Mukherjee, D., Gao, Y., Citri, A., Davison, I., & Mizrahi, A. (2017). Functional plasticity of odor representations during motherhood. Cell Reports, 21(2), 351–365. 10.1016/j.celrep.2017.09.038

Willott, J. F., Aitkin, L. M., & McFadden, S. L. (1993). Plasticity of auditory cortex associated with sensorineural hearing loss in adult C57BL/6J mice. Journal of Comparative Neurology, 329(3), 402–411. 10.1002/cne.903290310

Winkowski, D. E., & Kanold, P. O. (2013). Laminar transformation of frequency organization in auditory cortex. The Journal of Neuroscience, 33(4), 1498–1508. 10.1523/JNEUROSCI.3101-12.2013

Winters, C., Gorssen, W., Wöhr, M., & D’Hooge, R. (2023). BAMBI: A new method for automated assessment of bidirectional early-life interaction between maternal behavior and pup vocalization in mouse dam-pup dyads. Frontiers in Behavioral Neuroscience, 17. 10.3389/fnbeh.2023.1139254

Wong, G. T. (2002). Speed congenics: Applications for transgenic and knock-out mouse strains. Neuropeptides, 36(2), 230–236. 10.1054/npep.2002.0905

Zhang, Q., Liu, H., McGee, J., Walsh, E. J., Soukup, G. A., & He, D. Z. Z. (2013). Identifying microRNAs involved in degeneration of the organ of Corti during age-related hearing loss. PLOS ONE, 8(4), e62786. 10.1371/journal.pone.0062786

Zheng, Q. Y., Johnson, K. R., & Erway, L. C. (1999). Assessment of hearing in 80 inbred strains of mice by ABR threshold analyses. Hearing Research, 130(1), 94–107. 10.1016/S0378-5955(99)00003-9

